# Shared biosynthetic architectures generate diverse β-amino polyketide residues in cyanobacterial peptides

**DOI:** 10.64898/2026.06.02.729631

**Authors:** Aaditi Chopade, Anmol Chaure, Mary-Candler Schantz, Runjie Xia, David E. Berthold, Forrest W. Lefler, H. Dail Laughinghouse, Matthew J. Bertin

**Author notes:** Matthew J. Bertin, **Email:**. **Author Contributions:** M.J.B. and D.L. IV designed research; A.C., M-C.S., D.E.B., and F.W.L performed research; All authors analyzed data; M.J.B. wrote the first draft of the paper with input and approval from all authors. **Competing Interest Statement:** The authors declare no competing interests.

## Abstract

Cyanobacteria generate structurally complex peptides with diverse biological functions and vast chemical diversity, yet the enzymatic logic that produces their many documented unusual β-amino acid–containing polyketide units remains incompletely understood. Here we show that the recently described genus *Floridanema* harbors biosynthetic systems that unify the production of tychonamides and pahayokolide-like peptides, including the assembly of the signature residues 3-amino-2,5,7-trihydroxy-8-phenyloctanoic acid (Atpoa) and 3-amino-2,5,7,8-tetrahydroxy-10-methylundecanoic acid (Athmu). Using an integrated “pathways-to-products” strategy combining comparative genomics, bioinformatics analyses, and structure elucidation by NMR and MS, we link previously cryptic gene clusters to new metabolites and define shared architectural features across these pathways. Furthermore, we identified a ketoreductase in multiple pathways predicted to reduce α-keto acids and generate the starting unit in the Athmu moiety. These results support a common evolutionary origin or convergent phenomenon for constructing β-amino polyketide building blocks within cyanobacterial hybrid assembly lines. Together, our findings reposition *Floridanema* as a central source organism for these peptide families and establish biosynthetic principles that can be leveraged to predict, discover, and engineer related natural products.

**Significance Statement:** Cyanobacteria produce unusually complex peptides, but the enzymatic logic that builds their signature β-amino polyketide residues is poorly understood. By linking gene clusters identified in the genomes of members of the recently described *Floridanema* genus to the production of tychonamides, pahayokolide-like peptides, and other metabolites, we reveal shared biosynthetic architectures and suggest evolutionary strategies that drive peptide diversification.

## Introduction

Cyanobacteria are prolific producers of structurally diverse natural products, including many peptides and hybrid polyketide–peptide molecules assembled by nonribosomal peptide synthetases (NRPSs) and polyketide synthases (PKSs) that exhibit biosynthetic construction uncommon in other microorganisms (1). Among these metabolites are structurally complex “cyanopeptides” that incorporate nonproteinogenic amino acid residues and polyketide building blocks, a hallmark of NRPS and NRPS/PKS assembly-line biosynthesis (2). A recurring feature of multiple freshwater cyanobacterial peptide families is the presence of long-chain α-hydroxy-β-amino acid polyketide-derived residues, including Athmu (3-amino-2,5,7,8-tetrahydroxy-10-methylundecanoic acid) and Atpoa (3-amino-2,5,7-trihydroxy-8-phenyloctanoic acid) (3,4). These residues occur in structurally related cyclic peptides such as the pahayokolides (3), the portoamides (5), the lyngbyazothrins (6), the schizotrins (7), the puwainaphycins (8), the muscotoxins (9), and the tychonamides (4), and contribute substantially to molecular complexity, bioactivity, and diversification across these related molecules.

Pahayokolides A and B were originally isolated from a freshwater cyanobacterium identified as a *Lyngbya* sp. from the Florida Everglades and were shown to contain the unusual polyhydroxylated β-amino acid Athmu, consistent with a mixed NRPS/PKS biosynthetic origin (3). Follow-up stable-isotope incorporation experiments supported a biosynthetic model in which Athmu derives from a leucine-related starter unit that is extended by acetate units, with leucine providing the strongest isotopic enrichment and thus implicated as an immediate precursor to Athmu formation following conversion to α-ketoisocaproate (α-KIC) (10). These data are consistent with a pathway in which an NRPS loading module selects an α-keto acid (α-KIC) and subsequent on-assembly-line reduction produces an α-hydroxy functionality, and we provide evidence for this phenomenon in the current report, but our bioinformatic evidence points to the generation of α-KIC from isopropylmalate. This observation is similar to established paradigms for α-keto acid reduction in heterotrophic bacterial metabolites and related assembly lines (11,12).

Despite these insights, the enzymatic logic by which cyanobacterial hybrid NRPS/PKS systems repeatedly generate Athmu- and other β-amino polyketide building blocks remains incompletely resolved. More generally, genome-enabled natural product research has highlighted that many biosynthetic gene clusters (BGCs) remain “orphaned” (unlinked to products), and that comparative genomics paired with mass spectrometry can provide a route to connect cryptic BGCs to small-molecule structures and thereby reveal shared biosynthetic principles (13). For cyanobacterial systems in particular, elucidating how related pathways specify distinct long-chain β-amino polyketide residues—and how these pathways are distributed across taxa—would help explain both the origin of this chemistry and the evolutionary processes that drive peptide diversification.

Here we investigate biosynthetic systems from the recently described cyanobacteria *Floridanema* that unifies the production of tychonamides and pahayokolide/lyngbyazothrin/portoamide-like peptides, including the assembly of the hallmark β-amino polyketide residue Athmu. Using an integrated “pathways-to-products” strategy that combines comparative genomics and bioinformatics with structure elucidation by NMR and MS, we link gene clusters to new metabolites and identify shared architectural features across these pathways. In doing so, we provide evidence for conserved biosynthetic strategies for constructing β-amino polyketide building blocks within cyanobacterial hybrid assembly lines and reposition *Floridanema* as a key source organism for these peptide families.

## Results

### Genome mining links a hybrid PKS–NRPS gene cluster in *Floridanema flaviceps* to a new Athmu-containing peptide, floridanemamide (1)

Genome mining of the recently described *Floridanema* revealed multiple hybrid PKS–NRPS pathways predicted to produce large cyanopeptides containing unusual β-amino polyketide residues. Building on our earlier identification of the tychonamide pathway in *F. aerugineum* (14), we examined additional *Floridanema* genomes and identified in *F. flaviceps* (BLCC-F50) a closely related biosynthetic gene cluster (BGC), here named *fma*, that encodes a hybrid PKS–NRPS assembly line with tailoring domains consistent with biosynthesis of an Athmu-like β-amino polyketide residue. The putative pathway contained five open reading frames consisting of PKS architecture predicted to load a leucine-like substrate and to carry out three ketoextensions with amino and hydroxy modification of the final two acetate incorporations followed by the incorporation of 11 amino acids by NRPS modules (Fig. 1). LC–MS analysis of *F. flaviceps* extracts revealed an abundant metabolite with *m/z* 760.8985 [M+2H]^2+^, supporting the formula C_73_H_109_N_13_O_22_ (*SI Appendix*, Fig. S1). Guided by the predicted substrate specificities of the Fma NRPS adenylation domains (antiSMASH and PARAS reannotation) (15,16) and by diagnostic NMR correlations, we elucidated the structure of this compound as a new cyclic peptide–polyketide hybrid, which we name floridanemamide A (**1**) (Fig. 1). An examination of 1D and 2D NMR (*SI Appendix*, Fig. S2-S11A & Table S1) spectra suggested a large peptidic molecule and careful inspection of the HSQC-TOCSY spectrum allowed assignment of several amino acid residues which were also predicted from the bioinformatics analysis of the adenylation domains and tailoring enzymes in the BGC, which predicted an NRPS portion of D-Gln-Gly-Ser-Phe-D-Ile-Ser-Thr-Thr-D-NMe-Htyr-Pro-TE-Pro-TE. The dual thioesterase (TE) domains have been observed in other pathways such as the tychonamide pathway (14) and in that example, the final amino acid was linked to the PKS portion of the molecule. The BGC data were in almost complete harmony with the NMR data (Fig. 1). However, the first threonine residue predicted in the pathway was assigned as a dehydrobutyrine unit based on HSQC correlations between *δ*_C_ 120.6 and *δ*_H_ 5.70 (CH) and *δ*_C_ 13.1 and *δ*_H_ 1.81 (CH_3_) and correlations in the HMBC spectrum. The *N*-methyl homologated tyrosine was elucidated, not as an *N*-methyl amino acid, but as an *O*-methyl homologated tyrosine residue based on a TOCSY spin system *δ*_H_ 4.45 (CH), 1.88 CH_2_), and 2.54, 2.44 (CH_2_), doublet signals at *δ*_H_ 6.82 (CH) and *δ*_H_ 7.10, the HSQC correlation between *δ*_C_ 57.0 and *δ*_H_ 3.70 (CH_3_), and the HMBC correlation between *δ*_H_ 3.70 and *δ*_C_ 157.3. The *O*-methyl versus *N*-methyl annotation has been discussed previously in the putative biosynthesis of tychonamide and other specialized metabolites (14). Inferring configuration from the biosynthetic modules, this assigned the NRPS portion of the molecule as D-Gln-Gly-Ser-Phe-D-Ile-Ser-Dhb-Thr-D-*O*Me-Htyr-Pro with the final predicted proline not in the pure peptide portion of the molecule. The putative *fma* PKS region is organized into a short module set predicted to initiate from an α-keto acid–derived starter (α-ketoisocaproic acid) and to extend with malonyl units, followed by tailoring consistent with installation of both β-amino and polyhydroxylated functionalities. Notably, the cluster encodes an aminotransferase and monooxygenase domain combination also observed in the tychonamide pathway, supporting a conserved strategy for generating the 3-amino-2-hydroxy motif characteristic of Athmu-type residues. NMR (an extended TOCSY spin system) established the polyketide-derived unit in floridanemamide as Athmu (3-amino-2,5,7,8-tetrahydroxy-10-methylundecanoic acid) and placed it as the linkage point to the terminal proline in the BGC. The oxymethine proton attached to the fifth position in the unit was deshielded (*δ*_H_ 4.97) and NOE and HMBC correlations connected it to the final proline unit which was modified by an *N*-linked butyric acid moiety as supported by a TOCSY spin system connecting *δ*_H_ 2.24 (CH_2_), 1.48, (CH_2_), and 0.87 (CH_3_) with HMBC correlations to *δ*_C_ 171.7 (C=O). This modified proline residue has not been reported in a cyanobacterial metabolite but has been annotated in the fungal peptaibol culicinin D (17). This elucidation constructed the planar structure of floridanemamide A (**1**). We propose that the following D-amino acids are incorporated into the peptide macrocycle based on epimerization domains in the pathway (D-Gln & D-*O*Me-Htyr) and we confirmed that the predicted D-Ile residue was D-*allo*-Ile following derivatization of the amino acids in the peptide using a derivatizing agent (L-FDLA) and comparison to authentic amino acid standards similarly derivatized (*SI Appendix*, Fig. S12A). The D-*allo*-isoleucine is common in molecules of this class (3-6). We predict that the remaining amino acids are of the L-configuration. We next assigned the relative and absolute configuration of key centers in the Athmu residue, integrating bioinformatic predictions of ketoreductase (KR) stereochemistry with experimental data. Sequence motifs in the two β-ketoreducing KR domains predict alternating A- and B-type outcomes consistent with installation of defined stereocenters along the Athmu backbone (D- and L-products, respectively) (*SI Appendix*, Fig. S12B) (18). These predictions were supported by NMR-based 1,3-diol relationship analysis (19). Next chiral derivatization of the hydrolysate of **1** and Marfey-type analyses confirmed the configuration at C-3 (amino-bearing center) as *S* (20) (*SI Appendix*, Fig. S12C). The hydroxy bearing C-2 was assigned using *J*-coupling analysis (*SI Appendix*, Fig. S12D). A small coupling constant was determined for H_2_-H_3_ (2.9 Hz) following DQF-COSY analysis, and a small coupling constant was determined for H_2_-C_4_ (2.8 Hz) following inspection of the HETLOC spectrum. Finally, a small coupling constant was estimated for C_1_-H_3_ by the ratio of the relative magnitudes of cross peaks measured in the HMBC spectrum with respect to a common proton (21). This established a *threo* relation between C-2 and C-3. To assign the final configuration at C-8, *J*-coupling analysis and NOE correlations suggested a *threo* relationship between H-7 and H-8 with a medium value for ^3^*J*(H_2_-H_3_) = 5.2 Hz and small couplings for H_8_-C_6_ and H_7_-C_9_ (2.2 Hz and 3.5 Hz, respectively) (*SI Appendix*, Fig. S12D). We also turned to derivative formation and generated an acetonide derivative of **1** and the equivalence of the geminal dimethyls in the acetonide (*δ*_C_ 27.0) supported a *syn* configuration between C-7 and C-8 (22) (*SI Appendix*, S13). Together, these data assign Athmu in **1** as 2*R*,3*S*,5*R*,7*S*,8*S* and the peptide portion as D-Gln-Gly-Ser-Phe-D-*allo*-Ile-Ser-Dhb-Thr-D-*O*Me-Htyr-Pro, establishing floridanemamide A as a stereochemically defined Athmu-containing member of the *Floridanema* cyanopeptide class.

**Figure 1.**
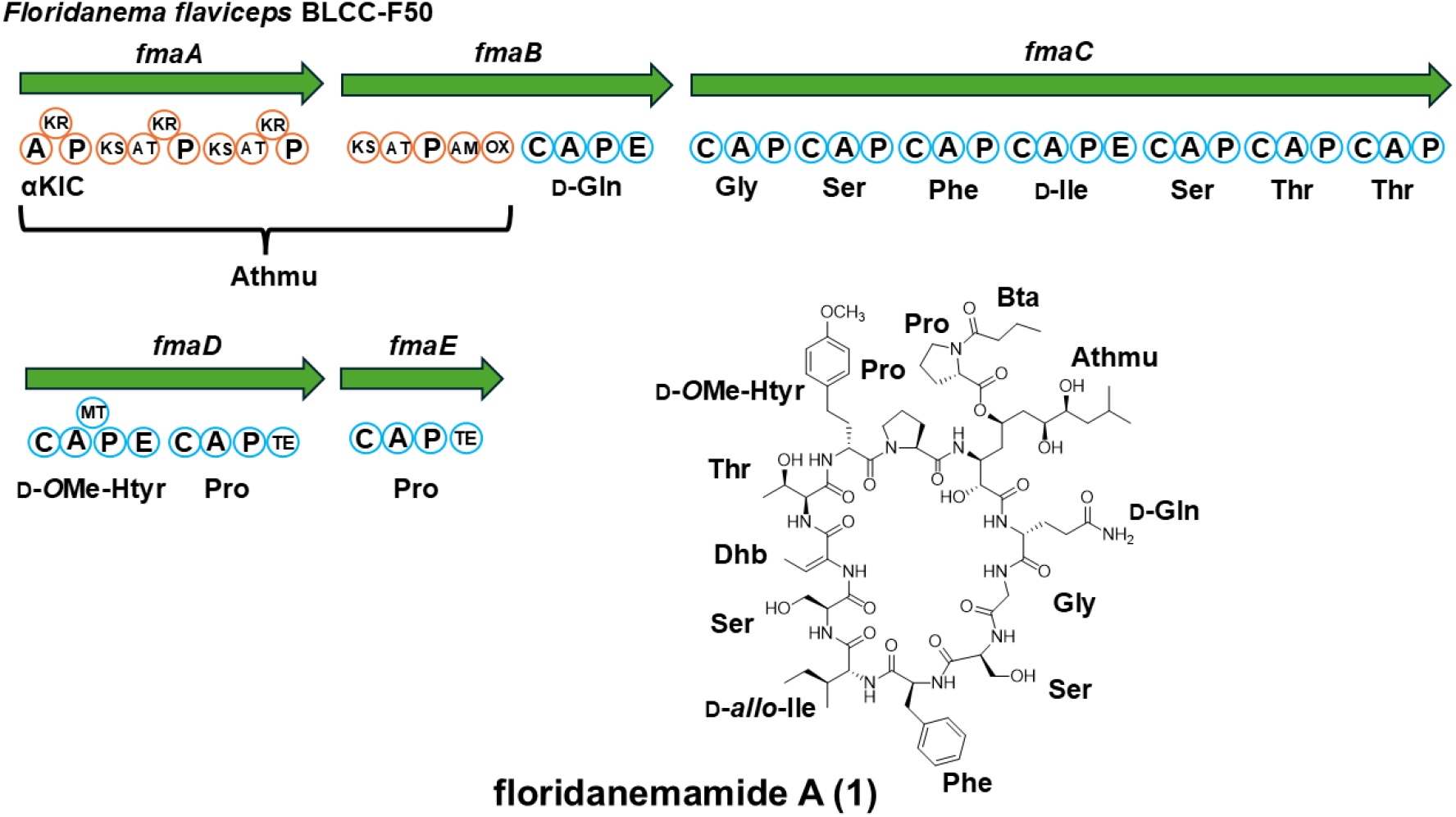
Organization of the putative floridanemamide biosynthetic pathway (*fma*) with open reading frames, modules, and domains annotated along with the floridanemamide structure. Amino acid predictions from antiSMASH and PARAS are below each module. Abbreviations: A, adenylation; P, carrier protein; KR, ketoreductase; KS, ketosynthase; AT, acyltransferase; AM, aminotransferase; OX, monooxygenase; C, condensation; E, epimerization; MT, methyltransferase; TE thioesterase.

### Bioinformatic evidence signals the biosynthetic construction of the α-hydroxyisocaproate starter unit

Stable-isotope feeding studies have implicated leucine-derived α-KIC as the starting unit in the Athmu residue, consistent with initiation from an α-keto acid followed by iterative malonyl extensions (10). In the putative *fma* biosynthetic pathway, we identified three adjacent enzymes annotated as 2-isopropylmalate synthase, 3-isopropylmalate dehydratase, and 3-isopropylmalate dehydrogenase, which together constitute a canonical route to α-KIC and suggest local biosynthesis of the α-keto acid starter (Fig. 2A). The loading-module ketoreductase (KR) contains conserved catalytic motifs consistent with an active reductase and clusters phylogenetically with KRs previously shown to reduce α-keto acid starter units in hybrid assembly lines (Fig. 2B). Taken together, these features support a biosynthetic model in which α-KIC is generated proximal to the assembly line and is reduced on assembly to α-hydroxyisocaproate (α-HIC) during the initiation of Athmu biosynthesis. This initiation logic mirrors strategies proposed for other Athmu-containing pathways such as the portoamides and lynbyazothrins. We identified the putative portoamide biosynthetic pathway in the genome of *Phormidium* sp. LEGE 05292 and a putative pahayokolide-like biosynthetic pathway from a *Floridanema* species (described below), which also both contained the isopropylmalate synthase, isopropylmalate dehydratase, and isopropylmalate dehydrogenase genes adjacent to the putative pathways and KR domains in the loading modules, which also both clustered with the KR from the *fma* pathway following phylogenetic analysis (Fig. 2B) further supporting the pathway to the starting unit.

**Figure 2.**
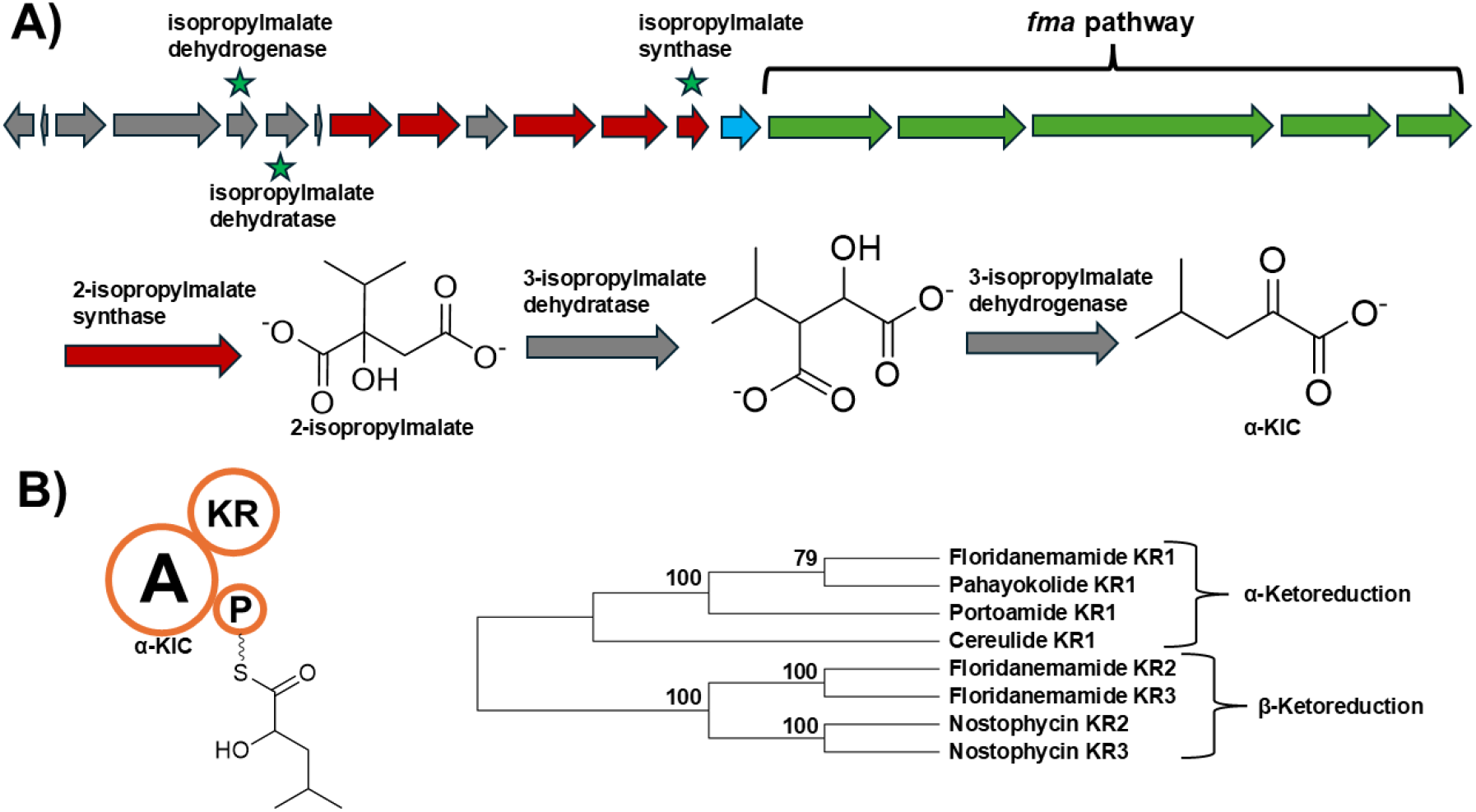
The generation of the α-hydroxyisocaproic acid starter unit in floridanemamide. (A) Organization of the contig from which the putative *fma* pathway was identified with the predicted genes for the synthesis of α-ketoisocaproic acid noted with green stars. The predicted biochemical transformations that create the α-KIC starting unit are shown below the pathway. (B) Organization of the Fma loading domain with a KR enzyme predicted to perform reduction at the α-ketone in α-KIC. A phylogenetic tree is organized at right showing the floridanemamide KR clustering with a KR from the cereulide pathway (Cer KR1), which was experimentally shown to reduce at the α-ketone position. The KR in the loading domain of the putative portoamide pathway also clusters with these KRs. For abbreviations, see Figure 1 legend.

### Connecting cyanopeptide chemical space and phylogenetic analysis repositions the *Floridanema* genus as a prolific producer of PKS-NRPS metabolites

Cyanobacterial natural products are frequently named for the initially assigned producer or collection site (e.g., *Tychonema*, tychonamide; *Lyngbya*, lyngbyazothrin; Porto, Portugal, portoamide), but ongoing taxonomic revisions can obscure relationships between metabolite families and their true producer lineages. To place floridanemamide A within the broader cyanopeptide landscape, we performed structure-based clustering of ∼3,000 cyanobacterial metabolites from CyanoMetDB (23) using SMILES-derived fingerprints and pairwise Tanimoto similarity, followed by dimensionality reduction and network visualization (Methods; Fig. 3A-C). This analysis positioned floridanemamide A within a tight cluster of highly related PKS–NRPS peptides, including lyngbyazothrins A–D, pahayokolides A and B, portoamides, tychonamides, and schizotrin A (Fig. 3A-C). Inspection of the original isolation reports for these compounds revealed recurring uncertainty or subsequent revision in the taxonomic assignment of the producing strains. For example, the pahayokolide producer showed the highest similarity of 93% to strains in the GenBank database based on 16S rRNA gene analysis (3), and the portoamide producer was reclassified from *Oscillatoria* sp. to *Phormidium* sp. (24). The lyngbyazothrin producer was reclassified from *Lyngbya* sp. SAG 36.91 to *Tenebriella* (25). Additionally, we isolated tychonamide A and new tychonamide analogs and identified the putative gene cluster for these metabolites in a strain of *Floridanema aerugineum* (14). However, the work from Moretto et al. (2024) showed strong phylogenetic evidence that the *Lyngbya*/*Tenebriella* SAG 36.91 should be reclassified as a *Floridanema* species (26). We therefore compiled available 16S rRNA sequences for reported producers (*Phormidium* sp. LEGE 05292, portoamides, GUO85101.1; Cyanobacterium PL, DQ072163.1 – the original tychonamide A producer) and reconstructed a phylogeny together with reference sequences used to define *Floridanema* (Fig. 4). In the resulting tree, sequences from strains reported to produce lyngbyazothrins, portoamides, and tychonamide-like peptides grouped within the *Floridanema* clade (Fig. 4), consistent with a shared producer lineage for these structurally related metabolites. Together with our identification of a tychonamide biosynthetic gene cluster and new tychonamide analogs from *Floridanema aerugineum*, these results support the conclusion that *Floridanema* represents a prolific and underrecognized source of this PKS–NRPS peptide family and that several historically attributed producers likely belong to *Floridanema* or closely related taxa. The widespread isolation of these strains from many freshwater communities, including the isolation of BLCC-F306 from Lake Erie, and the unique chemistry that this group produces provokes intriguing questions as to the role *Floridanema* plays in cyanobacterial community dynamics. Field sampling in July 2025 by our group in Lake Erie and subsequent MAG analysis from a cyanobacterial bloom field enrichment identified a partial gene cluster with nearly 100% amino acid identity to the tychonamide biosynthetic pathway from *Floridanema aerugineum* (*SI Appendix*, S14).

**Figure 3.**
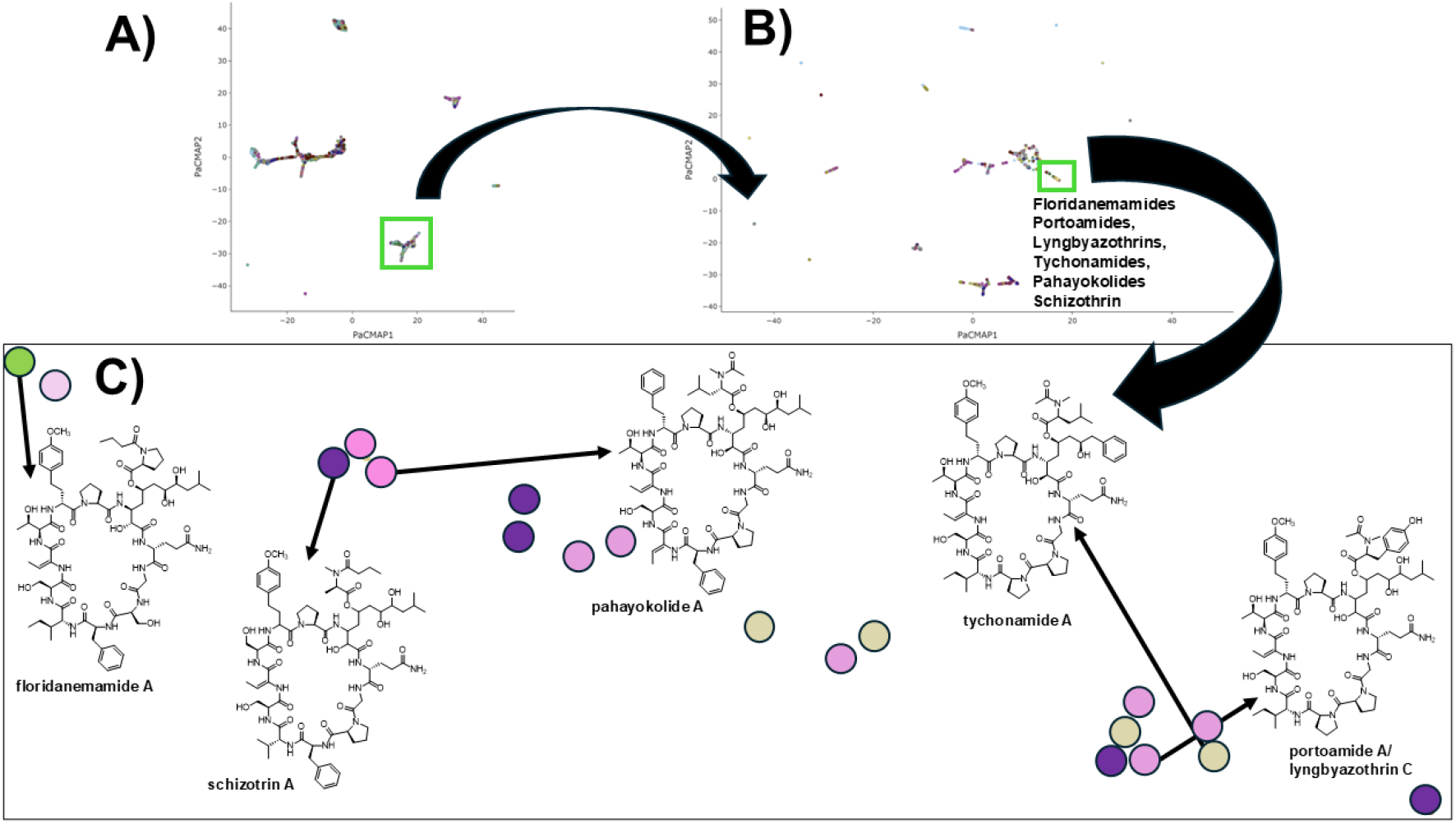
(A) Molecular network following the analysis of Tanimoto similarity of the CyanMetDB (t=0.30) with clustering via PaCMAP (n_neighbors = 10 and iterations = 20000). The green box shows a group of closely related compounds which were (B) subjected to an additional clustering step. (C) Enhanced cluster of these related compounds (green box in B) with representative structures in the network.

**Figure 4.**
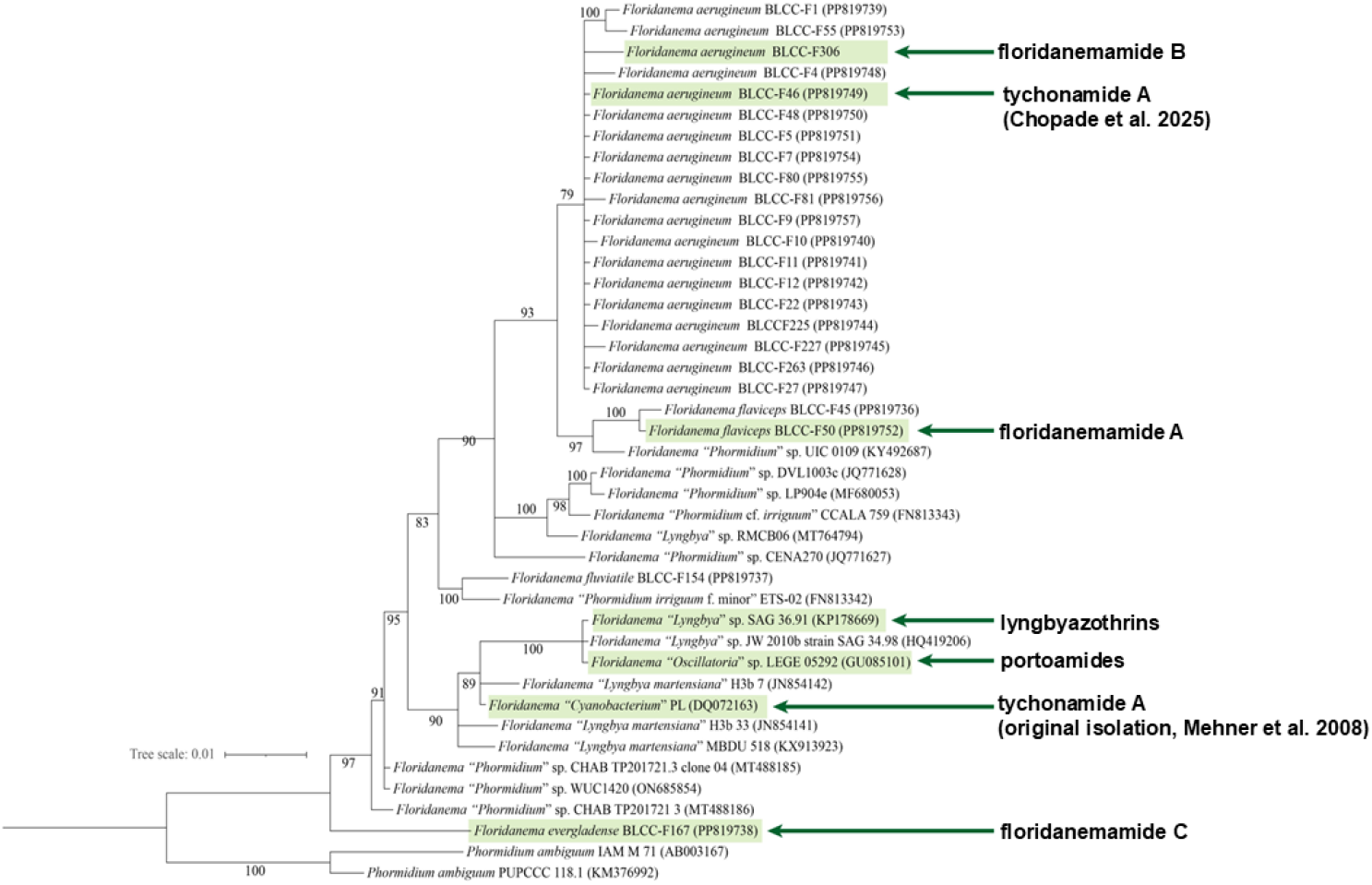
Phylogenetic analysis of 16S rRNA sequences from putative *Floridanema* strains with specialized metabolites noted for specific strains.

### Comparative genomics identifies a putative pahayokolide, portoamide, and additional floridanemamide biosynthetic pathways and supports at least three conserved initiation strategies that diversify β-amino polyketide residues in cyanobacterial PKS–NRPS peptides

To test whether Athmu- and Atpoa-class β-amino polyketide residues arise from shared biosynthetic architecture across taxa, we analyzed genomes from multiple *Floridanema* strains and representative publicly available cyanobacterial genomes associated with Athmu/Atpoa-containing metabolite families. This comparative analysis identified BGC architectures consistent with the biosynthesis of floridanemamide A-like/pahayokolide-like peptides in *Floridanema* sp. BLCC-F306 (*fmb* pathway) (Fig. 5A) and *Floridanema* sp. BLCC F167 (*fmc* pathway) (Fig. 5B) and portoamide-like peptides in “*Phormidium*” sp. LEGE 05292 (SAMN16351719) (*por* pathway) (*SI Appendix*, S15), with module order, tailoring domains, and predicted substrate selections matching the major structural features elucidated for these compounds (3,5). Although the modules in the putative portoamide pathway were split onto several contigs. Minor discrepancies (e.g., unresolved acetyltransferases in the *fmb* and *por* pathways and an unresolved *N*-methyltransferase and one apparently absent PKS extension in the portoamide-like pathway) are detailed in Fig. 5A and S14, but do not alter the shared logic of the loading and early extension steps that define the β-amino residue class. Additionally, in *fmbD* we would predict the incorporation of a likely *O*Me-Htyr residue like the other documented peptides in this class, but pahayokolides A and B incorporate Hphe in this position (3), possibly indicating some flexibility in substrate selection or a mutation in this adenylation domain in the *Floridanema* strain analyzed. Tychonamide B incorporates the Hphe residue, while tychonamide A has an *O*Me-Htyr further supporting this flexibility (4). However, it could be that a *Floridanema* strain contains a full biosynthetic gene cluster with an adenylation domain that prefers homologated phenylalanine. We were able to isolate and elucidate the structure of the putative product from *Floridanema aerugineum* BLCC-F306, which had a high-resolution mass measurement of *m/z* 751.8934 [M+2H]^2+^ supporting a formula of C_73_H_107_N_13_O_21_ and the NMR data were in near complete harmony with the putative biosynthetic pathway predictions including the incorporation of two Dhb units (from Thr) and the *O*Me-Htyr unit and we named this compound floridanemamide B (**2**) (Fig. 4A and *SI Appendix*, Table S2 and Fig S16-S23). We also found a low abundance molecular feature in the BLCC-F306 extract with *m/z* 736.8872 [M+2H]^2+^ supporting a molecular formula of C_72_H_105_N_13_O_20_, which is consistent with pahayokolide A (calcd *m/z* 736.8872, C_72_H_107_N_13_O_20_^2+^) (*SI Appendix*, S24). We were also able to isolate a polyketide-peptide from the *Floridanema evergladense* BLCC-F167 extract (*m/z* 757.9400 [M+2H]^2+^ supporting a formula of C_73_H_119_N_13_O_21_) (SI Appendix Fig S25), which again showed near complete harmony between the predicted biosynthetic pathway and the structure itself (elucidated from 1D and 2D NMR data), which we named floridanemamide C (**3**) (Fig. 5B and *SI Appendix*, Table S3 and Fig 11B and Fig S26-S32). This new metabolite had the same butylated proline residue seen in **1**, and the Marfey’s analysis demonstrated that the Ile residue was D-*allo*-Ile as in **1** (*SI Appendix*, S12A). Interestingly, the absolute configuration of the 2-hydroxy-3-amino functionality was of the opposite configuration in **2** and **3** (*S,R*) compared to **1** (*R,S*), determined by Marfey’s analysis and *J*-coupling. Inspection of KR sequences and additional *J*-coupling analysis completed the remaining absolute configuration of the Athmu units in **2** and **3** (*SI Appendix*, S12 B-D), which was the same as that of **1**. Predicted protein annotations for all biosynthetic pathways discussed in this report can be found in SI Appendix, Table S4.

**Figure 5.**
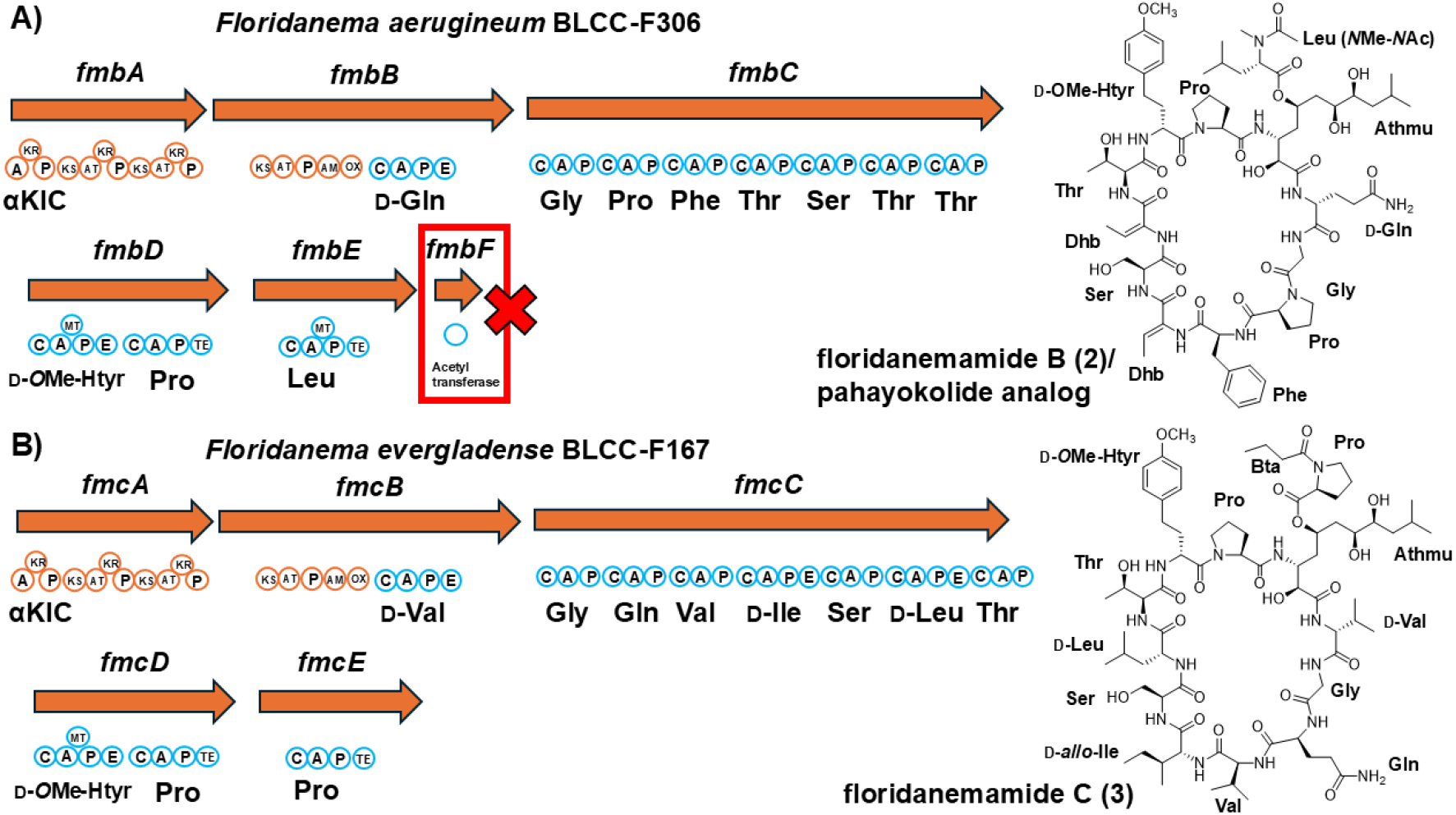
Putative floridanemamide B gene cluster (*fmb*) and floridanemamide C (*fmc*) biosynthetic pathways with structures shown at right. Red boxes show portions of pathways that were not annotated but are predicted for biosynthesis. For abbreviations, see Figure 1 legend.

Comparing gene organization with product composition across PKS–NRPS cyanopeptides that contain 3-amino-2-hydroxy β-amino acid residues reveals patterns of diversification. Pathway similarity analysis identifies *fmaC* and *fmaE* as the most divergent modules (Fig. 6A). Consistent with this, the corresponding amino acid positions in the peptide products show the greatest variability (Fig. 56; SI Appendix, Table S5). Together, these results indicate that sequence diversification is concentrated in specific modules and the β-amino acid unit, suggesting a mechanism for tuning peptide structure—and potentially target engagement and ecological function—through localized changes in amino acid incorporation.

**Figure 6.**
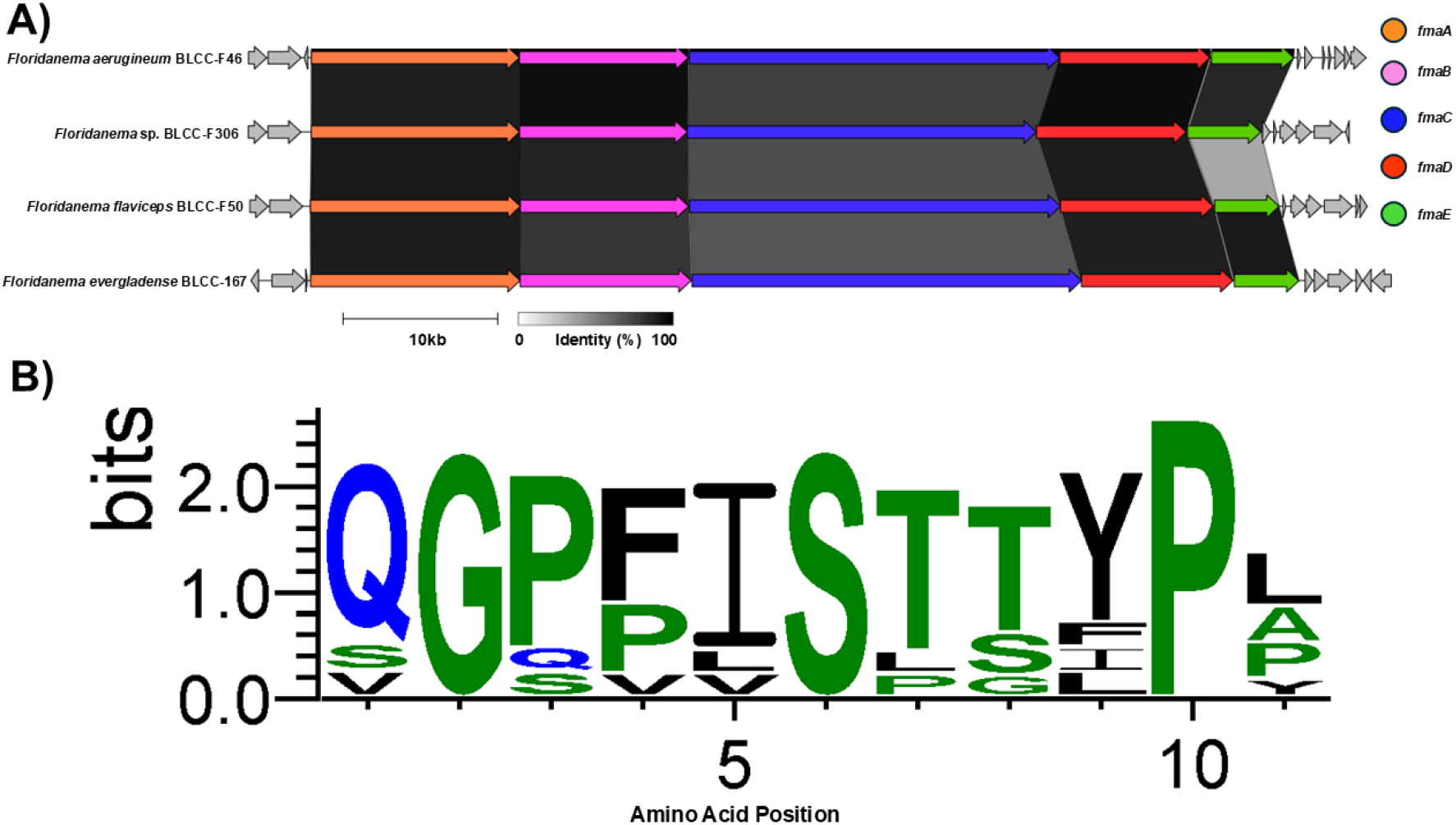
(A) Synteny plot of the open reading frames (based on the *fma* pathway) for the putative pathway of tychonamides (BLCC-F46) and 1-3. (B) Consensus amino acid sequences for PKS-NRPS molecules with 3-amino-2-hydroxy beta amino acids and ten amino acid peptide cyclic portions: tychonamide A, floridanemamides A-C, pahayokolide A, portoamide A/lyngbyazothrin C, schizotrin A, scytonemin A, and muscotoxin A. The consensus sequences are noted with proteinogenic amino acids, but often amino acids are modified (e.g., *N*-methyl homologated tyrosine at position 9). The alignment was created using WebLogo v3.9.0 (https://weblogo.threeplusone.com/). The full sequences for compounds can be found in Table S4.

The Comparison across Athmu- and Atpoa-containing pathways revealed that diversification of 3-amino-2-hydroxy–bearing β-amino polyketide residues is concentrated in the initiation module (“starting unit”) and can be grouped into three recurring strategies (Fig. 7A-C). First, Athmu-class residues are predicted to initiate from an α-keto acid selected by an adenylation domain (e.g., α-ketoisocaproate) followed by on-assembly-line reduction to the corresponding α-hydroxy starter, a logic shared by floridanemamides, pahayokolide-like, and lyngbyazothrin/portoamide-like pathways (Fig. 7A). Second, Atpoa/Ahoa/Ahda-class residues are associated with a CoA-ligase (Cal) initiation that recruits an aromatic acid starter (e.g., phenylacetate) and is followed by distinct PKS module arrangements and late-stage amination/oxygenation, consistent with the structural divergence observed among these residue types (e.g., tychonamide (14), scytonemin A (27), and nostophycin (28) (Fig. 7B). Third, Cal-type initiation without an embedded reductase is linked to recruitment of fatty-acyl starters that yield long-chain β-amino residues such as Ahmos in puwainaphycin/minutissamide-like pathways (29-32) (Fig. 7C).

**Figure 7.**
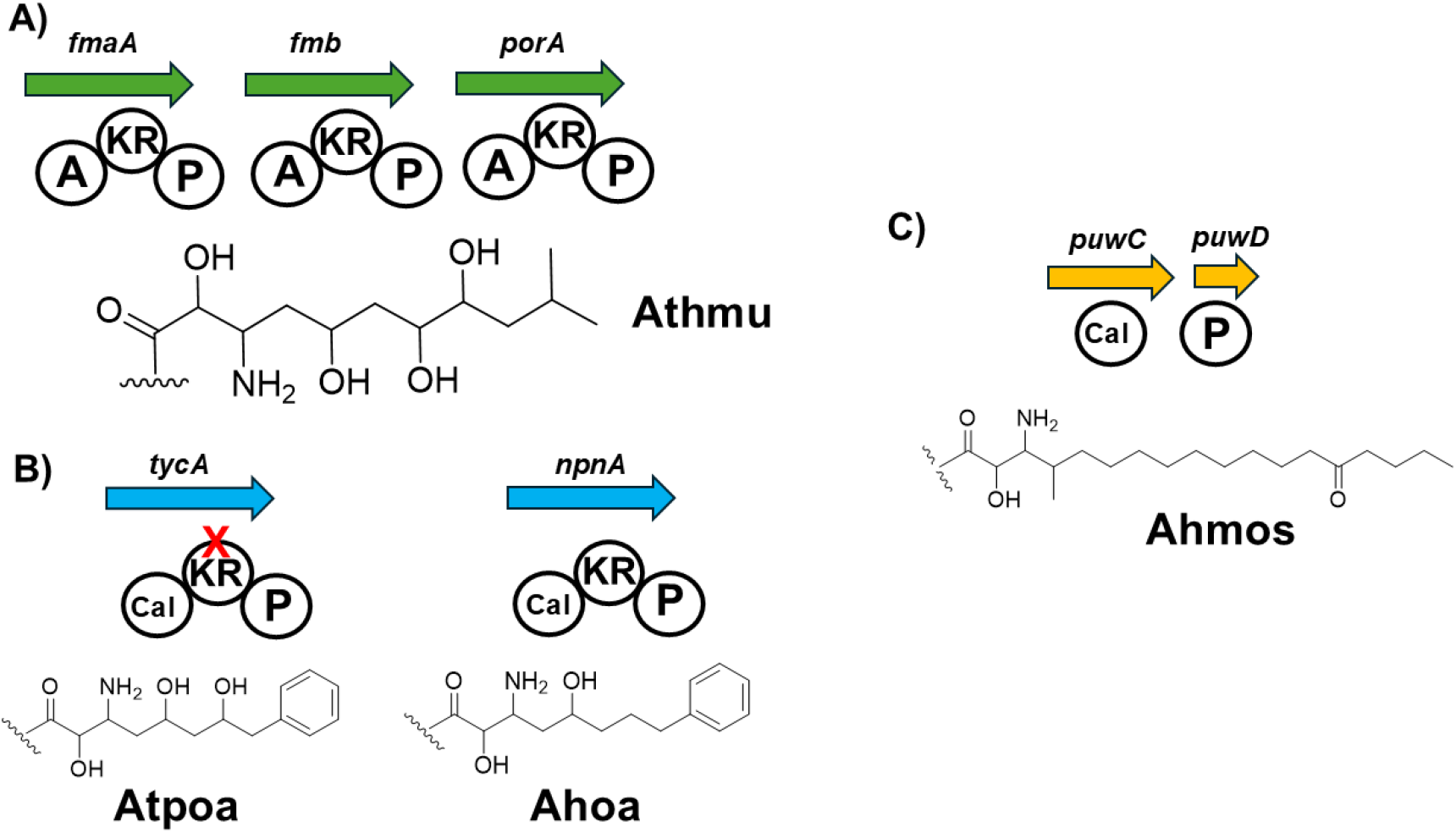
The substrate loading domains for α-hydroxy-β-amino acid polyketide-derived residues. (A) The putative floridanemamide and portoamide initiation modules that ultimately generate the Athmu unit. (B) The putative tychonamide and nostophycin initiation modules that ultimately generate Atpoa and Ahoa, respectively (KR domain of *tyc* is predicted to be inactive). (C) The putative initiation module for the puwainaphycins ultimately generating the Ahmos unit. For enzyme domain abbreviations, see Figure 1 legend.

## Discussion

Cyanobacterial hybrid PKS–NRPS pathways generate many structurally elaborate peptide natural products, yet the biosynthetic logic that repeatedly yields long-chain, polyoxygenated β-amino polyketide residues remains only partially understood. Collectively, the *fma*, and other pathways and the elucidated structures in this report extend the emerging view that members of *Floridanema* harbor a set of related hybrid assembly lines that generate Athmu/Atpoa-class β-amino polyketide building blocks using shared enzymatic assemblies, providing a mechanistic foundation for diversification across tychonamide- and floridanemamide/pahayokolide-like peptide families.

Our results support a key mechanistic insight in that Athmu biosynthesis can be programmed by α-keto acid initiation coupled to on-assembly reduction. Feeding studies previously implicated leucine-derived α-ketoisocaproate (α-KIC) as a precursor to Athmu (10). Conversely, our data strongly support a conserved genomic signature consistent with a new model: isopropylmalate pathway enzymes located adjacent to *fma* and to additional Athmu-associated clusters (portoamide), together with an active loading-module ketoreductase (KR) that groups phylogenetically with α-keto-reducing KRs. This combination supports a biosynthetic model in which α-KIC is locally constructed and reduced to α-hydroxyisocaproate (α-HIC) during initiation, linking primary metabolism to specialized metabolite starter-unit selection. It may be that the feeding of leucine and deamination and conversion to α-KIC occurred during previous feeding experiments and was incorporated but that leucine is not the true biogenic origin of the starting unit. However, the architectural adjacency argument alone cannot exclude scenarios in which the BGC-proximal isopropylmalate genes are not the functionally dominant source of α-KIC *in vivo* and future biochemical experiments can validate our hypothesis. There is precedence for the on-assembly reduction prediction in the generation of the cyanobacterial metabolites spumigin and certain aeruginosides (33,34) in which a keto acid (4-hydroxyphenylpyruvic acid) is reduced to 4-hydroxyphenyllactic acid in the loading domain. Future biochemical studies of the loading adenylation domain and KR function can elucidate substrate scope and function.

Together, these results position *Floridanema* as a central and underrecognized lineage for Athmu/Atpoa-bearing cyanopeptides and provides new generalizable principles for predicting related chemistry from genome sequence. Although, as 16S rRNA analysis can be insufficient to resolve closely related cyanobacterial lineages, additional phylogenomics will further test these relationships to understand the phylogeny of the producing taxa. By integrating structure-based clustering of ∼3,000 cyanobacterial metabolites with phylogenetic reassessment of reported producers, we provide evidence that multiple historically attributed peptide families likely originate from *Floridanema* or closely related taxa. Cyanobacterial natural products are often named for an initially assigned genus or collection site, but frequent taxonomic revisions—combined with limited sequence data in early isolation studies—can obscure producer lineages and fragment our understanding of pathway evolution. The proximity of floridanemamide to lyngbyazothrins, pahayokolides, portoamides, and tychonamides in chemical similarity space, together with 16*S*-based clustering of multiple “producer” strains within or near the *Floridanema* clade, supports a unifying producer framework for this PKS–NRPS peptide family (cf. Fig. 3A-C and Fig. 4). There are other effective chemical space organizing tools such as NPAtlas (35), and searching the floridanemamide structure with a similarity threshold of 0.6 identified a similar group of related molecules to our clustering tool: lyngbyazothrins A-D/portoamides A-D, muscotoxin A/C, pahayokolide A/B, schizotrin A, and tychonamide A/B. In the future, we will integrate phylogenomics and biosynthetic gene cluster information, and biogeography with our clustering and networking tool to investigate chemical diversity and biosynthetic evolution questions. The widespread presence of *Floridanema* in freshwater benthic cyanobacterial communities, and our detection of *Floridanema* in a Lake Erie cyanoHAB, combined with evidence that related metabolites exhibit cytotoxic and allelopathic activities (4,5), suggests that this lineage may influence freshwater cyanobacterial community structure through chemically mediated interactions. These observations motivate future work to quantify *Floridanema* occurrence across bloom stages, determine how its metabolite production affects competitor taxa and grazers, and assess whether these compounds contribute meaningfully to exposure-relevant bloom chemistry and public health risk.

We also organize Athmu/Atpoa diversification into a small set of recurring “starter-unit programming” strategies. More broadly, comparative genomics supports at least three conserved initiation strategies sufficient to explain much of the β-amino polyketide residue diversity in cyanobacterial peptides: (1) α-keto acid selection by an adenylation domain with loading-module reduction (Athmu-class), (2) aromatic-acid initiation by a CoA ligase (Atpoa/Ahoa/Ahda-class), and (3) fatty-acyl initiation by a CoA ligase yielding long-chain β-amino residues (Ahmos/Hamd/Hamh-class). This framework suggests that residue-level diversification is driven primarily by starter programming and early tailoring, rather than wholesale pathway rewiring. However, we did not perform a quantitative census across all available cyanobacterial genomes (instead relying on a chemistry-first approach) and estimates of prevalence and conservation (and potential exceptions) will benefit from a broader, systematic survey and targeted validations, which we will pursue. Remaining discrepancies (e.g., unresolved acetyltransferase assignments; apparent amino acid substrate swaps in individual modules) define tractable targets for biochemical validation in future experiments. Collectively, these principles provide a roadmap to prioritize orphan clusters, anticipate β-amino residue classes, and ultimately leverage modular starter programming to diversify cyanobacterial peptide scaffolds, which will be useful in bioengineering and drug discovery. On the engineering side, the modularity of starter unit incorporation suggests that swapping or redesigning loading modules and early tailoring cassettes may offer an efficient route to generate new β-amino residue variants and thereby diversify peptide scaffolds. Retroevolutionary analysis may provide a blueprint to determine the shared ancestry and divergence events between the loading domains in the pathways discussed in this report and others that generate β-amino acids as well as the adenylation domain mutations that generate substrate flexibility in the NRPS portions of these molecules (36,37).

## Materials and Methods

### General experimental procedures

NMR spectra were recorded on a Bruker 500 MHz Ascend Avance III NMR instrument equipped with a multinuclear broadband cryoprobe, and the chemical shifts reported for the floridanemamides were referenced to the residual solvent peak of (CD_3_)_2_SO (Cambridge Isotope Laboratories) (*δ*_H_ 2.50 and *δ*_C_ 39.5). LC-HRMS and LC-HRMS^2^ data were collected on an Agilent Revident Q-TOF system equipped with a Jet Stream source and 1290 Infinity II Bio LC (with multisampler and multicolumn thermostat) and MassHunter Workstation software. Low resolution mass spectrometry data were obtained using an Agilent LC-MSD single quadrupole mass spectrometer equipped with an Agilent 1260 HPLC system including autosampler. Semipreparative HPLC was carried out using an Agilent 1260 system equipped with a vacuum degasser, autosampler, and diode array detector.

### Isolation and cultivation of cyanobacterial strains and the extraction and isolation of the floridanemamides

BLCC-F50 was originally isolated from Port Saint Lucie, FL, USA and BLCC-F167 was originally isolated from Everglades National Park, Homestead, FL, USA (26). BLCC was isolated from Lake Erie near The Stone Laboratory, Put-in-Bay, OH, USA. Uni-cyanobacterial cultures of *F. flaviceps* (strain: BLCC-F50), *F. evergladense* (strain: BLCC-F167), and *F. aerugineum* (strain: BLCC-F306) were cultured in replicate 2 L bioreactors using BG-11 medium and incubated at 25°C on a 12:12–h light: dark cycle under cool white (6500K) fluorescent lighting (15 µmol.photons.m-2.s-1) at the Berthold Laughinghouse Culture Collection (BLCC) (University of Florida-IFAS, Davie, FL, USA). Cyanobacterial cultures were harvested after 1 month of cultivation, frozen at -80°C, and lyophilized. The lyophilized biomass samples were shipped to the Bertin Laboratory at Case Western Reserve University (Cleveland, OH, USA) (BLCC-F50: 1.83 g dry weight; BLCC-F167: 1.19 g dry weight; BLCC F306: 1.07 g dry weight) and the samples were each repeatedly extracted with 100% CH_3_OH - each extraction stirred with gentle heat (30°C) for 30 min. The extracts of each specimen were combined and concentrated under reduced pressure (BLCC-F50: 430 mg; BLCC-F167: 119 mg; BLCC-F306: 113 mg). Next, the extracts were separated over a 10 g or 2 g C18 SPE cartridge eluting with CH_3_CN, which resulted in 93 mg of the CH_3_CN fraction for BLCC-F50 (BLCC-F167: 11.4 mg; BLCC-F306: 15.8 mg). The BLCC-F50 CH_3_CN fraction was subjected to semi-preparative HPLC using a Kinetex C18 5 µm column (250 mm × 10 mm) and an isocratic solvent system of 50% H_2_O in CH_3_CN with 0.1% formic acid added. Floridanemamide A (**1**) was purified from repeated chromatographic isolation (t_R_ = 5.8 min, 13.4 mg). The BLCC-F306 and BLCC-F167 CH_3_CN fractions were separated using the same isocratic method and solvent system used to isolate **1**, ultimately isolating **2** (t_R_ = 5.8 min, 3.3 mg) and **3** (t_R_ = 11.8 min, 2.2 mg).

*Floridanemamide A* (**1**): colorless oil; [*α*]^20^_D_ = +9.3 (*c* 0.1, MeOH); UV λ_max_: 210, 238, 278 nm (from HPLC-DAD); ^1^H and ^13^C NMR, see Table S1; HRESIMS *m/z* 760.8985 [M+H]^2+^ (calcd for C_73_H_111_N_13_O_22_^2+^, 760.8978).

*Floridanemamide B* (**2**): colorless oil; [*α*]^20^_D_ = -23 (*c* 0.033, MeOH); UV λ_max_: 210, 238, 278 nm (from HPLC-DAD); ^1^H and ^13^C NMR, see Table S2; HRESIMS *m/z* 751.8934 [M+H]^2+^ (calcd for C_73_H_109_N_13_O_21_^2+^, 751.8925).

*Floridanemamide C* (**3**): colorless oil; [*α*]^20^_D_ = +29 (*c* 0.025, MeOH); UV λ_max_: 210, 238, 278 nm (from HPLC-DAD); ^1^H and ^13^C NMR, see Table S3; HRESIMS *m/z* 757.9400 [M+H]^2+^ (calcd for C_73_H_121_N_13_O_21_^2+^, 757.9395).

### Lake Erie field sampling, DNA isolation, and MAG analysis

Surface water samples were collected in western Lake Erie on July 23, 2025, from Showse Park, Vermilion, OH, USA, 41.43014°N, 82.31420° during a cyanoHAB event. Approximately 1 L of surface water was collected into opaque amber high-density polyethylene (HDPE) bottles and transported to the laboratory on ice. Upon arrival, 60 mL aliquots were filtered through 47 mm, 0.2 μm Sterlitech polyester track etch (PETE) membrane filters and the two replicates were stored at –80°C for DNA extraction and molecular analyses. Next, laboratory enrichments were established in the laboratory by inoculating 1 mL of field water into 9 mL of BG-11 medium, with two replicate cultures. Cultures were maintained at 22°C under a 12 h light/12 h dark photoperiod for three weeks under 4100K LED lights (15 µmol•photons•m^−2^•s^−1^) which were kept 22 cm from the cultures. Biomass was collected by filtration through 47 mm, 0.2 μm membrane filters. Genomic DNA was extracted from one 10 mL laboratory culture using the Qiagen DNeasy Plant Mini Kit with minor modifications. Cells were pelleted by centrifugation at 4000 × g for 10 min, resuspended in 400 μL of AP1 buffer with 4 μL RNase A, and disrupted by bead beating for 1 min. Following lysis, DNA was purified and eluted in 200 μL of Buffer AE according to the manufacturer’s protocol. Concentration and purity were assessed using a NanoDrop 2000 spectrophotometer and a Qubit fluorometer prior to sequencing. Purified DNA was sequenced by Plasmidsaurus using the Oxford Nanopore Technologies platform and assembled by Plasmidsaurus into contigs using metaSPADES (38), which were analyzed via the bioinformatics pipeline described below. We divided the contigs into those over 1,000 bp (1,182 contigs; mean length 3805 bp; std. dev. 3453 bp; max length 35,178 bp) and those under 1000 bp (98 contigs; mean length 722 bp; std. dev. 142 bp; max length 993 bp).

### Marfey’s analysis

D-Ile, D-*allo*-Ile, L-Ile, and L-*allo*-Ile standards (MedChemExpress) were derivatized with N-α-(2,4-dinitro-5-fluorophenyl)-L-leucinamide (L-FDLA; TCI) by mixing each amino acid in 1 M NaHCO_3_(100 µL) with 1% L-FDLA in acetone and heating at 40 °C for 1 h. Reactions were quenched with 2 N HCl (50 µL). Floridanemamides A and C (**1 & 3**) (0.2 mg) were hydrolyzed in 6 N HCl (90 °C, 16 h), dried under N_2_, reconstituted in 1 M NaHCO_3_(100 µL), and derivatized with 1% L-FDLA as above. FDLA derivatives of standards and the hydrolysates were analyzed by LC–MS using two methods (Luna 5 μm C18 column: 150 mm × 2 mm; 0.4 mL/min; both mobile phases contained 0.1% formic acid): *Method* **1**, 20–80% H_2_O–CH_3_CN over 30 min (re-equilibration 31–36 min), separated the L- and D-series and supported the residue in **1** being in the D-series, but did not resolve Ile from *allo*-Ile within each series. *Method 2*, 25% H_2_O–CH_3_CN over 30 min using a YMC 5 μm Chiral ART Cellulose-SB column (250 mm × 4.6 mm), resolved D-Ile and D-*allo*-Ile and assigned the isoleucine residue in floridanemamides A and C as D-*allo*-Ile (Table 1).

**Table 1.** Retention times of Ile-L-FLDA derivatives using two different methods.

| Amino acid | t<sub>R</sub> gradient method 1 | t<sub>R</sub> isocratic method 2 |
| --- | --- | --- |
| L-isoleucine | 17.73 | - |
| L-*allo*-isoleucine | 17.65 | - |
| D-isoleucine | 21.88 | 9.95 |
| D-*allo*-isoleucine | 21.89 | 9.22 |
| Flordanemamide A hydrolysate | 21.88 | 9.19 |
| Flordanemamide C hydrolysate |  | 9.40 |

To determine the configuration of the amino bearing carbon in the Athmu units of **1-3**, a portion of each floridanemamide (0.7 mg) was also separately hydrolyzed using 6 N HCl and heated at 90°C for 16 h. The hydrolysates were divided into two equal portions and all were dried under a stream of N_2_, reconstituted in 100 μL of 1 M NaHCO_3_ solution, and derivatized with 500 μL of a 1% solution of N-α-(2,4-dinitro-5-fluorophenyl)-L-leucinamide (L-FDLA) (TCI America) or N-α-(2,4-dinitro-5-fluorophenyl)-D-leucinamide (D-FDLA) (TCI America) in acetone, followed by stirring and heating at 40°C for 1 h. The reaction mixtures were then cooled to room temperature and quenched with 50 μL of 2 N HCl. The hydrolysates were subsequently diluted 1:10 with 1:1 H_2_O/CH_3_CN to a final volume of 1 mL and analyzed by LC-MS using a Kinetex C18 column (150 mm × 4.6 mm) for analyte separation with a column heater temperature of 30°C. The separation was performed with a gradient of CH_3_CN in H_2_O from 30% to 70% over 40 min, with a flow rate of 0.2 mL/min with respect to **1** and 20% to 80% over 40 min, with a flow rate of 0.2 mL/min for **2** and **3**. The L-FDLA derivative eluted at 32.2 min and the D-FDLA derivative eluted at 40.4 min designating an *S* configuration at C-3 in floridanemamide A (**1**), while we saw the D-FDLA derivative elute before the L-FDLA derivative for **2** and **3**, designating an *R* configuration.

### Floridanemamide acetonide formation

Pure floridanemamide A (**1**, 0.5 mg) was added to a solution of p-toluenesulfonic acid (pTsOH) (2.27mg) and 2,2-dimethylpropane (0.6 mL) and stirred at room temperature for 24 h. The reaction was quenched with sodium bicarbonate (NaHCO_3_). The product was partitioned between ethyl acetate (EtOAc) and water. The resulting organic layer was dried under nitrogen gas and analyzed using NMR and HRMS. *Floridanemamide acetonide*: ^1^H NMR *SI Appendix*, S13; HRESIMS *m/z* 780.9140 [M+H]^2+^ (calcd for C_76_H_115_N_13_O_22_^2+^, 780.9134).

### Genome sequencing and assembly of cultivated specimens

Genomic libraries for Illumina sequencing were prepared using the Illumina TruSeq Library Construction Kit (Illumina, Inc., USA), and 2×150 bp paired-end reads were generated with Illumina NovaSeq (Illumina, Inc., USA) at UF ICBR (University of Florida’s Interdisciplinary Center for Biotechnology Research) NextGen DNA Sequencing Core Facility (RRID:SCR_019152). Genomic libraries for Oxford Nanopore Technologies (ONT) sequencing were prepared using the Technologies Native Barcoding Kit (#SQK-NBD114) and the Long Fragment Buffer was used to promote longer read lengths. Sequencing was performed on the PromethION 2 Solo platform using a FLO-PRO114M Flow Cell (R10 version). Trimming was done using fastp v1.0.1 (39) and Barbell v0.3.1 (40) for Illumina and ONT, respectively. Assembly of ONT reads was done using myloasm v0.2.0 (41), polished using Dorado v1.3.0 (Oxford Nanopore Technologies; https://github.com/nanoporetech/dorado) and PYPOLCA (42) for ONT and Illumina reads, respectively. Circular contigs were identified and the remaining contigs were binned using COMEBin v1.0.4 (43), MetaDecoder v1.2.0 (44), SemiBin2 v2.1.0 (45), taxVAMB v5.0.3 (46) and further refined with Binette v1.1.2 (47). Final bin quality was assessed using CheckM2 v1.0.1 (48), coverage was determined using CoverM v0.7.0 (49), taxonomy was assigned using GTDB-tk (R226) v2.4.1 (50), annotated with Bakta v1.11.4 (51). Summaries are available in *SI Appendix*, Table S6.

### Bioinformatics and phylogenetic analysis

The assembled genomes of *F. flaviceps* BLCC-F50, *F. evergladense* BLCC-F167, and *F. aerugineum* BLCC-F306, sequenced via Oxford Nanopore and Illumina HiSeq (26), were publicly available (BioSample #s: SAMN43549591, SAMN43549593 and SAMN60208900, respectively). The assembled genomes were uploaded to antiSMASH (version 8) (15) with the Extra Features tab set at “All on”. Hybrid biosynthetic gene clusters were analyzed for modules and domains consistent with those predicted to generate hybrid polyketide-peptide molecules. Additionally, the genome of “*Phormidium*” sp. LEGE 05292 (SAMN16351719) was publicly available, and it was analyzed in the same manner as the *Floridanema* genomes. The Lake Erie MAG sample was analyzed for contigs with GC content between 40%-50%, which were subjected to BLASTp analysis within the Geneious Prime 2019 software platform and a contig of interest with multiple NRPS domains (alignment hit to *Anabaena* sp.) was uploaded to antiSMASH as described above where reanalysis using the BLASTp function embedded within antiSMASH showed 100% identity to *F. aerugineum* (100% identity, 1795 accession length). Sequences for the nostophycin (*npn*, BGC0001029.5) and puwainaphycin A (*puw*, BGC0001125.5) gene clusters were accessed from MIBiG (52).

Phylogenetic analyses of the 16 rRNA sequences included closely related taxa that were identified using BLAST (Basic Local Alignment Search Tool, NCBI). Multiple sequences alignments (MSAs) from Moretto et al., (2024) (26) and MSA’s for *Floridanema* were retrieved from CyanoSeq V1.3 (53). The MSAs were aligned using MUSCLE5 (54) and trimmed with trimAl v1.5.0 (55) and the best fit substitution model was determined with ModelTest-NG v0.1.7 (56) using corrected Akaike Information Criterion. Maximum likelihood (ML) analysis was carried out using RaxML-NG v1.2.2 (57). Bayesian inference (BI) was conducted with MrBayes using 1.0 × 10^6^ generation, a 0.25 burn–in rate, and resampling every 100 generations, (58). The synteny plots were visualized using clinker v0.0.32 (59).

### Cheminformatics and networking generation

SMILES strings were extracted for all molecules in the CyanoMetDB (23), a database containing 3085 cyanobacterial specialized metabolites. Before the analysis, the database was pre-processed by standardizing inconsistent label variations, removing invalid SMILE chains, and merging duplicate entries. The chemical structures were then converted into fingerprints for further processing using the RDKit’s Morgan fingerprint function (60), which encodes the local chemical environment of each atom into fixed-length binary vectors, and has an established noise threshold based on the distribution of similarities between randomly chosen molecules. A pairwise Tanimoto similarity matrix was then computed from the resulting Morgan2 fingerprints. Upon inspection of the similarity distribution, it was revealed that the majority of pairwise similarity scores fell below 0.2, reflecting the broad structural diversity of the database. Therefore, to retain only strongly correlated pairs, an adaptive threshold was applied based on statistical methods, and all entries below this value were set to zero. For dimensionality reduction, PCA and PaCMAP were sequentially applied, as PCA alone was unable to preserve the dataset’s properties (61). Furthermore, PaCMAP was chosen for its robust preservation of both local and global structure across the dataset. The resulting outputs were visualized interactively using Plotly (62), revealing five major clusters within the chemical space. While this provided a generalized overview of the chemical space, a more detailed analysis of the local structure within the floridanemamides cluster was required. Additionally, the previously used adaptive threshold had condensed the space by discarding weak similarity pairs, reducing the local resolution further. To recover this local structure for the cluster of floridanemamides, the entire pipeline was repeated independently. The corresponding metadata was extracted, and a new pairwise Tanimoto similarity matrix was computed for 566 molecules. In this second pass, the adaptive threshold was also omitted, as applying a sparsity-inducing cutoff in this context would have eliminated meaningful chemical relationships rather than uninformative noise. The resulting embedding yields a higher-resolution view of the local chemical space within this structurally cohesive group.

## Data and Software Availability

Genome assemblies are available for *F. flaviceps* BLCC-F50 (SAMN43549591), *F. evergladense* BLCC-F167 (SAMN43549593), and *F. aerugineum* BLCC F306 (SAMN60208900) on NCBI GenBank. NMR data for the floridanemamides were deposited at NP-MRD and will be available following publication (A: NP0354017; B: NP0354018; C: NP0354019). Code is available at https://github.com/anmolchaure/AC_AC-cyanoMet-cheminformatics-.

## Acknowledgments

Research reported in this article was supported by the National Institute of Environmental Health Sciences of the National Institutes of Health under award number R21ES033758 (M. J. B.) and the National Institute of Food and Agriculture, U.S. Department of Agriculture, Hatch project award number FLA-FTL-006614 (H. D. L.). The content is solely the responsibility of the authors and does not necessarily represent the official views of the National Institutes of Health.

## Notes

### Competing Interest Statement

The authors have declared no competing interest.

