## Supporting Information for "Shared biosynthetic architectures generate diverse β-amino polyketide residues in cyanobacterial peptides"

**This PDF file includes:**

Table S1-S6
Figures S1-S32

Table S1. NMR data for floridanemamide A (**1**) (500 MHz for $^1$H and 125 MHz for $^{13}$C, DMSO-$d_6$)

| Unit | No | $\delta_C$, multi. | $\delta_H$ [mult., $J$ (Hz)] | HMBC | TOCSY |
|---|---|---|---|---|---|
| Pro-1 | 1 | 171.4, C | | | |
| | 2 | 55.7, CH | 4.27, ovlp | 1 | 2, 3a, 3b, 5b |
| | 3a | 27.8, $CH_2$ | 1.98, m | | 2, 3b, 5b |
| | 3b | | 1.72, m | | 3a, 3b, 5b |
| | 4 | 23.9, $CH_2$ | 1.85, ovlp | | 2 |
| | 5a | 46.7, $CH_2$ | 3.50, m | | 2, 3a, 3b |
| | 5b | | 3.33, m | | 3b |
| D-*O*-me-Htyr | 6 | 169.6, C | | | |
| | 7 | 50.4, CH | 4.45, m | 6 | NH, 8, 9a, 9b |
| | 8 | 33.0, $CH_2$ | 1.88, m | 9 | NH, 7, 9a, 9b |
| | 9a | 30.2, $CH_2$ | 2.54, m | 8 | NH, 7, 8, 9b |
| | 9b | | 2.44, m | | NH, 7, 8 |
| | 10 | 133.4, C | | | |
| | 11 | 129.2, CH | 7.10, d, 8.5 | 10, 13 | 12 |
| | 12 | 113.7, CH | 6.82, d, 8.5 | 10, 13 | 11 |
| | 13 | 157.3, C | | | |
| | 14 | 113.7, CH | 6.82, d, 8.5 | 10, 13 | 15 |
| | 15 | 129.2, CH | 7.10, d, 8.5 | 10, 13 | 14 |
| | 16 | 55.0, $CH_3$ | 3.70, s | 13 | |
| | NH | | 7.93, m | | 7, 8, 9a, 9b |
| Thr | 17 | 169.9, C | | | |
| | 18 | 58.9, CH | 4.35, ovlp | 17 | NH, 20 |
| | 19 | 66.8, CH | 4.22, m | | 20 |
| | 20 | 20.2, $CH_3$ | 1.07, d, 6.3 | 19 | NH, 18, 19 |
| | NH | | 7.82, m | | 18, 19, 20 |
| Dhb | 21 | 163.9, C | | 23 | |
| | 22 | 131.5, C | | 23 | |
| | 23 | 120.6, CH | 5.70, m | 21 | 24 |
| | 24 | 13.1, $CH_3$ | 1.81, d, 7.3 | 23 | NH |
| | NH | | 9.35, m | | 23, 24 |
| Ser-1 | 25 | 170.5, C | | | |
| | 26 | 56.6, CH | 4.16, m | 25 | NH, 27a, 27b |
| | 27a | 61.8, $CH_2$ | 3.69, m | 26 | NH |
| | 27b | | 3.55, m | | NH |
| | NH | | 8.14, m | | 26, 27a, 27b |
| D-*allo*-Ile | 28 | 171.0, C | | | |
| | 29 | 58.5, CH | 4.47, m | 28 | NH, 30 |
| | 30 | 35.3, CH | 1.85, ovlp | 33 | NH, 32, 33 |
| | 31 | 25.3, $CH_2$ | 1.00, m | 33 | 32, 33 |
| | 32 | 11.6, $CH_3$ | 0.74, t, 7.3 | 31 | NH, 30, 31 |

| | | | | | |
|---|---|---|---|---|---|
| | 33 | 14.5, $CH_3$ | 0.69, d, 6.4 | 31 | |
| | NH | | 8.60, m | | 29, 30, 31, 32, 33 |
| Phe | 34 | 172.0 | | | |
| | 35 | 54.7, CH | 4.50, m | 34 | NH, 36a, 36b |
| | 36a | 36.5, $CH_2$ | 3.01, m | 35 | NH, 35, 36a |
| | 36b | | 2.97, m | | |
| | 37 | 138.4, C | | | |
| | 38 | 129.3, CH | 7.23, m | 37 | |
| | 39 | 128.2, CH | 7.27, m | | |
| | 40 | 126.7, CH | 7.24, m | 39, 41 | |
| | 41 | 128.2, CH | 7.27, m | | |
| | 42 | 129.3, CH | 7.23, m | 37, 41 | |
| | NH | | 8.11, m | | 35, 36 |
| Ser-2 | 43 | 170.7, C | | | |
| | 44 | 55.7, CH | 4.32, ovlp | 43 | NH, 45 a |
| | 45a | 61.3, $CH_2$ | 3.72, m | | NH, 44 |
| | 45b | | 3.57, m | | NH, 44 |
| | NH | | 8.12, m | | 44, 45b |
| Gly | 46 | 170.3, C | | | |
| | 47 | 41.7, $CH_2$ | 3.78, m | 46 | NH |
| | NH | | 8.20, m | | 47 |
| D-Gln | 48 | 172.0, C | | | |
| | 49 | 52.4, CH | 4.24, ovlp | 48 | NH, 50, 51 |
| | 50 | 28.7, $CH_2$ | 1.92, m | 51 | NH, 49, 51 |
| | 51 | 31.5, $CH_2$ | 2.09, m | 52 | NH, 49,50 |
| | 52 | 174.9, C | | | |
| | $NH_2$ | n.o.[a] | | | |
| | NH | | 7.95, m | | 49, 50, 51 |
| Athmu | 53 | 171.5, C | | | |
| | 54 | 72.0, CH | 3.94, m | 53 | NH, 56a |
| | 55 | 48.7, CH | 4.09, m | | NH, 54, 56 a, 56b, 57, 59 |
| | NH | | 7.60, m | | 54, 55, 56a, 56b, 57, 58 |
| | 56a | 37.9, $CH_2$ | 1.77, ovlp | | NH, 55, 58, 60 |
| | 56b | | 1.43, m | | NH, 55, 58, 60, 61, 63 |
| | 57 | 69.9, CH | 4.97, m | | NH, 55, 56a, 56b, 58, 59 |
| | 58 | 38.0, $CH_2$ | 1.47, m | | NH, 54, 55, 56a, 56b, 60, 61, 63, 64 |

| | | | | | |
|---|---|---|---|---|---|
| | 59 | 71.2, CH | 3.20, m | | 57, 58, 61, 62, 63, 64 |
| | 60 | 71.5, CH | 3.29, m | 59 | 57, 58, 61, 62, 63, 64 |
| | 61 | 41.7, $CH_2$ | 1.19, m | 62 | 60, 62, 63, 64 |
| | 62 | 23.5, CH | 1.75, ovlp | 61 | NH, 55, 58, 60, 61, 63 |
| | 63 | 23.9, $CH_3$ | 0.86, ovlp | 62 | 58, 60, 61, 62, 64 |
| | 64 | 21.7, $CH_3$ | 0.81, d, 6.6 | 62 | 63 |
| Pro-2 | 65 | 172.2 | | 57, 66 | |
| | 66 | 58.4, CH | 4.46 m | 67 | 67a |
| | 67a | 28.6, $CH_2$ | 1.93, m | | 66 |
| | 67b | | 1.79, ovlp | | 67a |
| | 68 | 21.7, $CH_2$ | 1.75, ovlp | | 67a |
| | 69a | 46.0, $CH_2$ | 3.46, m | | 67a, 68 |
| | 69b | | 3.29, m | | 67b, 68 |
| Butyric acid | 70 | 171.7, C | | | |
| | 71 | 35.7, $CH_2$ | 2.24, m | 70, 72 | 72, 73 |
| | 72 | 18.1, $CH_2$ | 1.48, m | 70, 73 | 71, 73 |
| | 73 | 13.6, $CH_3$ | 0.87, ovlp | | 71, 72 |

[a]not observed

Table S2. NMR data for floridanemamide B (**2**) (500 MHz for $^{1}H$ and 125 MHz for $^{13}C$, DMSO-$d_6$)

| Unit | No | $\delta_c$, multi. | $\delta_H$ [mult., $J$ (Hz)] |
|---|---|---|---|
| Pro-1 | 1 | 167.0, C | |
| | 2 | 60.2, CH | 4.27, m |
| | 3 | 28.0, $CH_2$ | 1.93, m |
| | 4 | 24.8, $CH_2$ | 1.86, m |
| | 5a | 46.7, $CH_2$ | 3.48, m |
| | 5b | | 3.34, m |
| D-*O*-me-Htyr | 6 | 166.9, C | |
| | 7 | 51.5, CH | 4.32, m |
| | 8a | 33.4, $CH_2$ | 1.87, m |
| | 8b | | 1.70, m |
| | 9a | 30.7, $CH_2$ | 2.55, m |
| | 9b | | 2.41, m |
| | 10 | 133.8, C | |
| | 11 | 129.5, CH | 7.10, d, 8.3 |
| | 12 | 113.7, CH | 6.82, d, 8.3 |
| | 13 | 157.5, C | |
| | 14 | 113.7, CH | 6.82, d, 8.3 |
| | 15 | 129.5, CH | 7.10, d, 8.3 |
| | 16 | 55.3, $CH_3$ | 3.70, s |
| | NH | | 7.97, m |
| Thr | 17 | n.o.[a] | |
| | 18 | 59.8, CH | 4.26, m |
| | 19 | 66.9, CH | 4.14, m |
| | 20 | 20.1, $CH_3$ | 1.08, d, 6.3 |
| | NH | | 7.48, m |
| Dhb-1 | 21 | 164.3 | |
| | 22 | 141.2 | |
| | 23 | 124.4, CH | 5.64, q, 7.2 |
| | 24 | 13.2, $CH_3$ | 1.83, s |
| | NH | | 9.28, m |
| Ser-1 | 25 | 172.4, C | |
| | 26 | 56.5, CH | 4.22, m |
| | 27a | 61.4, $CH_2$ | 3.77, m |
| | 27b | | 3.69, m |
| | NH | | 8.00, m |
| Dhb-2 | 28 | 164.9, C | |
| | 29 | 140.0, C | |
| | 30 | 129.7, CH | 5.76, m |
| | 31 | 13.2, $CH_3$ | 1.83, s |
| | NH | | 9.02, m |
| Phe | 32 | 170.6, C | |

| | | | |
|---|---|---|---|
| | 33 | 56.48, CH | 4.35, m |
| | 34a | 36.7, $CH_2$ | 3.04, m |
| | 34b | | 1.61, m |
| | 35 | 137.5, C | |
| | 36 | 129.0, CH | 7.29, ovlp |
| | 37 | 129.0, CH | 7.29, ovlp |
| | 38 | 126.7, CH | 7.23, m |
| | 39 | 129.0, CH | 7.29, ovlp |
| | 40 | 129.0, CH | 7.29, ovlp |
| | NH | | 7.99, m |
| Pro-2 | 41 | 172.8, C | |
| | 42 | 58.8, CH | 4.41, m |
| | 43 | 28.2, $CH_2$ | 1.88, m |
| | 44 | 21.8, $CH_2$ | 1.66, m |
| | 45 | 46.0, $CH_2$ | 3.47, m |
| Gly | 46 | 168.5, C | |
| | 47 | 41.2, $CH_2$ | 3.87, m |
| | NH | | 7.95, m |
| D-Gln | 48 | 170.2, C | |
| | 49 | 52.8, CH | 4.20, m |
| | 50 | 28.4, $CH_2$ | 1.92, m |
| | 51 | 31.5, $CH_2$ | 2.11, m |
| | 52 | 174.5, C | |
| | $NH_2$ | | n.o. |
| | NH | | 7.67, m |
| Athmu | 53 | 171.0, C | |
| | 54 | 72.0, CH | 3.86, m |
| | 55 | 48.6, CH | 4.00, m |
| | NH | | 7.56, m |
| | 56 | 36.7, $CH_2$ | 1.76, m |
| | 57 | 69.6, CH | 5.03, m |
| | 58 | 37.7, $CH_2$ | 1.41, m |
| | 59 | 70.9, CH | 3.14, m |
| | 60 | 72.0, CH | 3.27, m |
| | 61 | 41.9, $CH_2$ | 1.18, m |
| | 62 | 23.9, CH | 1.71, m |
| | 63 | 21.6, $CH_3$ | 0.83, ovlp |
| | 64 | 23.4, $CH_3$ | 0.88, ovlp |
| *N*-Me-Leu | 65 | 170.5, C | |
| | 66 | 53.7, CH | 5.08, m |
| | 67 | 36.3, $CH_2$ | 1.62, m |
| | 68 | 24.3, CH | 1.38, m |
| | 69 | 21.3, $CH_3$ | 0.87, ovlp |

| | | | |
|---|---|---|---|
| | 70 | 23.2, $CH_3$ | 0.89, ovlp |
| | 71 | 32.0, $CH_3$ | 2.82, s |
| Ac | 72 | 170.7. C | |
| | 73 | 21.6, $CH_3$ | 2.00, s |

[a]not observed

Table S3. NMR data for floridanemamide C (**3**) (500 MHz for $^1$H and 125 MHz for $^{13}$C, DMSO-$d_6$)

| Unit | No | $\delta_C$, multi. | $\delta_H$ [mult., $J$ (Hz)] | HMBC | TOCSY |
|---|---|---|---|---|---|
| Pro-1 | 1 | | | | |
| | 2 | 58.5, CH | 4.25, m | | 3, 4, 5a |
| | 3 | 28.4, $CH_2$ | 2.06, m | | 5a |
| | 4 | 24.3, $CH_2$ | 1.87, ovlp | | 2, 5a |
| | 5a | 46.9, $CH_2$ | 3.47, m | | 2, 3, 4 |
| | 5b | | 3.34, m | | |
| D-*O*-me-Htyr | 6 | | | | |
| | 7 | 51.2, CH | 4.45, m | | 8, 9a, NH |
| | 8 | 33.9, $CH_2$ | 1.88, m | | 7, 9a, NH |
| | 9a | 30.9, $CH_2$ | 2.56, m | 8, 10 | 7, 8, NH |
| | 9b | | 2.50, m | | |
| | 10 | 134.7, C | | | |
| | 11 | 129.3, CH | 7.10, d, 8.3 | 13 | 12 |
| | 12 | 113.6, CH | 6.81, d, 8.3 | 10 | 11 |
| | 13 | 157.9, C | | | |
| | 14 | 113.6, CH | 6.81, d, 8.3 | 10 | 15 |
| | 15 | 129.3, CH | 7.10, d, 8.3 | 13 | 14 |
| | 16 | 55.3, $CH_3$ | 3.71, s | 13 | |
| | NH | | 8.01, ovlp | | 7 |
| Thr | 17 | | | | |
| | 18 | 57.6, CH | 4.39, m | | 19, 20, NH |
| | 19 | 67.2, CH | 4.04, m | | 20, NH |
| | 20 | 19.9, $CH_3$ | 1.04, d, 6.0 | 19 | 18, 19 |
| | NH | | 8.17, m | | 18, 19 |
| Leu | 21 | | | | |
| | 22 | 55.3, CH | 4.46, m | | NH, 23, 24, 25, 26 |
| | 23 | 38.3 $CH_2$ | 1.78, m | | NH, 22, 24, 25, 26 |
| | 24 | 24.1, CH | 1.56, m | | 22, 23, 25, 26 |
| | 25 | 19.4, $CH_3$ | 0.85, ovlp | 24 | 22, 23, 24 |
| | 26 | 19.2, $CH_3$ | 0.79, ovlp | 24 | 22, 23, 24 |
| | NH | | 8.00, ovlp | | 22, 23 |
| Ser-1 | 27 | | | | |
| | 28 | 56.3, CH | 4.28, m | | 29a, 29b, NH |
| | 29a | 62.4, $CH_2$ | 3.65, m | | 28, NH |
| | 29b | | 3.58, m | | 28, NH |
| | NH | | 8.01, ovlp | | 28, 29a, 29b |
| D-*allo*-Ile | 30 | | | | |
| | 31 | 58.6, CH | 4.47, m | | 33, 34, 35, NH |
| | 32 | 36.3, CH | 1.97, m | | 33, 34, 25 |

| | | | | | |
|---|---|---|---|---|---|
| | 33 | 25.9, $CH_2$ | 1.12, m | | 32, 34, 35 |
| | 34 | 11.6, $CH_3$ | 0.82, t, 7.0 | 33 | 32, 33 |
| | 35 | 14.7, $CH_3$ | 0.79, d, 7.2 | 33 | 32, 33 |
| | NH | | 8.30, m | | 31 |
| Val-1 | 36 | 170.2, C | | | |
| | 37 | 59.5, CH | 4.11, m | | NH, 38, 39, 40 |
| | 38 | 30.5, CH | 1.98, ovlp | | NH, 37, 39, 40 |
| | 39 | 19.0, $CH_3$ | 0.88, d, 6.7 | 38 | 37, 38 |
| | 40 | 18.8, $CH_3$ | 0.92, d, 6.7 | 38 | 37, 38 |
| | NH | | 8.23, m | | 37, 38, 39, 40 |
| Gln | 41 | | | | |
| | 42 | 52.3, CH | 4.24, m | | NH, 43, 44 |
| | 43 | 28.5, $CH_2$ | 1.88, ovlp | | NH, 42, 44 |
| | 44 | 32.0, $CH_2$ | 2.09, m | 43, 44 | NH, 42, 43 |
| | 45 | 174.9, C | | | |
| | NH | | 7.95, s | | 42 |
| | $NH_2$ | | 8.01, br | 45 | |
| Gly | 46 | 168.5, C | | | |
| | 47 | 41.7, $CH_2$ | 3.72, m | | NH |
| | NH | | 8.22, m | | 47 |
| D-Val-2 | 48 | 170.2, C | | | |
| | 49 | 58.8, CH | 3.98, m | | NH, 50, 51, 52 |
| | 50 | 36.2, CH | 1.98, ovlp | | NH, 49, 51, 52 |
| | 51 | 19.0, $CH_3$ | 0.87, ovlp | | 49, 50 |
| | 52 | 19.0, $CH_3$ | 0.88, ovlp | 50 | 49, 50 |
| | NH | | 7.79, m | 50 | 49, 50, 51, 52 |
| Athmu | 53 | 171.2, C | | | |
| | 54 | 73.2, CH | 3.97, m | | NH, 56, 59, 60 |
| | 55 | 48.8, CH | 4.05, m | | NH, 56, 57, 59, 60 |
| | NH | | 7.43, s | | NH, 55, 56 |
| | 56 | 37.9, $CH_2$ | 1.78, ovlp | | NH, 57, 59, 60, 61 |
| | 57 | 69.6, CH | 5.03, m | | NH, 55, 56, 58 |
| | 58 | 38.0, $CH_2$ | 1.41, m | | 57, 59, 62 |
| | 59 | 70.5, CH | 3.26, ovlp | | 58, 61, 62, 63, 64 |
| | 60 | 71.6, CH | 3.29, ovlp | | 61, 62, 63, 64 |
| | 61 | 41.9, $CH_2$ | 1.20, m | 62 | 60, 62, 63, 64 |
| | 62 | 24.0, CH | 1.76, ovlp | | 60, 61, 63, 64 |
| | 63 | 21.6, $CH_3$ | 0.82, ovlp | 62 | 62 |
| | 64 | 24.0, $CH_3$ | 0.86, ovlp | 62 | 62 |
| Pro-2 | 65 | 170.5, C | | | |
| | 66 | 53.7, CH | 4.47, m | | 67, 68, 69a |

| | | | | | |
|---|---|---|---|---|---|
| | 67 | 28.0, $CH_2$ | 1.85, m | | 66, 69, 69b |
| | 68 | 22.7 $CH_2$ | 1.75, m | | 66, 67, 69a, 69b |
| | 69a | 46.9 $CH_2$ | 3.49, ovlp | | 66, 67, 68 |
| | 69b | | 3.31, ovlp | | 68 |
| Butyric acid | 70 | 172.8, C | | | |
| | 71 | 35.5, $CH_2$ | 2.22, m | | 72, 73 |
| | 72 | 17.7, $CH_2$ | 1.50, m | 71, 73 | 71, 73 |
| | 73 | 13.9, $CH_3$ | 0.86, m | 72 | 71, 72 |

Table S4. Predicted Protein Annotations for *fma*, *fmb*, and *fmc* Biosynthetic Pathways

| ORF | Module | Size (nt) | Proposed Domain Organization/ Function (AntiSMASH) | Predicted Substrate | Similar Sequence | Identity (%) | Coverage (%) | E-value | Accession number |
|---|---|---|---|---|---|---|---|---|---|
| **Floridanemamide A (BLCC F50)** | | | | | | | | | |
| ***fma*A** | Loading | 13,587 | A, KR, ACP | α-ketoisocaproic acid | *Microseira wollei]* | 71.61 | 99 | 0 | WP_226584072.1 |
| | 1 | | KS, AT, KR, ACP | malonyl CoA | | | | | |
| | 2 | | KS, AT, KR, ACP | malonyl CoA | | | | | |
| ***fma*B** | 1 | 10,890 | KS, AT, AmT, MO | malonyl CoA | *Desmonostoc muscorum* CCALA 125 | 72.0 | 100 | 0 | UBH04462.1 |
| | 2 | | C, A, E, PCP | D-Gln | | | | | |
| ***fma*C** | 1 | 24,054 | C, A, PCP | Gly | *Desmonostoc muscorum* CCALA 125 | 60.3 | 100 | 0 | UBH04463.1 |
| | 2 | | C, A, PCP | L-Ser | | | | | |
| | 3 | | C, A, PCP | L-Phe | | | | | |
| | 4 | | C, A, E, PCP | D-*allo*-Ile | | | | | |
| | 5 | | C, A, PCP | L-Ser | | | | | |
| | 6 | | C, A, PCP | L-Thr | | | | | |
| | 7 | | C, A, PCP | L-Thr | | | | | |
| ***fma*D** | 1 | 9,900 | C, A, E, oMT, PCP | D-hTyr | *Desmonostoc muscorum* LEGE 12446 | 67.02 | 89 | 0 | WP_255264360.1 |
| | 2 | | C, A, PCP | L-Pro | | | | | |
| ***fma*E** | 1 | 4173 | C, A, PCP, TE | L-Pro | *Nostoc* sp. | 59.97 | 99 | 0 | WP_335055746.1 |
| **Floridanemamide B (BLCC F306)** | | | | | | | | | |
| ***fmb*A** | Loading | 13,485 | A, KR, ACP | α-ketoisocaproic acid | *Microseira wollei]* | 71.49 | 99 | 0 | WP_226584072.1 |
| | 1 | | KS, AT, KR, ACP | malonyl CoA | | | | | |
| | 2 | | KS, AT, KR, ACP | malonyl CoA | | | | | |
| ***fmb*Bp** | 1 | 10,857 | KS, AT, AmT, MO | malonyl CoA | *Desmonostoc muscorum* CCALA 125 | 72.60 | 100 | 0 | UBH04462.1 |
| | 2 | | C, A, E, PCP | D-Gln | | | | | |
| ***fmb*C** | 1 | 22,584 | C, A, PCP | Gly | *Desmonostoc muscorum* CCALA 125 | 63.47 | 100 | 0 | UBH04462.1 |
| | 2 | | C, A, PCP | L-Pro | | | | | |
| | 3 | | C, A, PCP | L-Phe | | | | | |
| | 4 | | C, A, E, PCP | L-Thr | | | | | |
| | 5 | | C, A, PCP | L-Ser | | | | | |
| | 6 | | C, A, PCP | L-Thr | | | | | |
| | 7 | | C, A, PCP | L-Thr | | | | | |
| ***fmb*D** | 1 | 9,687 | C, A, E, oMT, PCP | D-hTyr | *Desmonostoc muscorum* LEGE 12446 | 68.43 | 89 | 0 | WP_255264360.1 |
| | 2 | | C, C, A, PCP | L-Pro | | | | | |
| ***fmb*E** | 1 | 4,743 | C, A, nMT, PCP, TE | L-Leu | *Nostoc* sp. UHCC 0251 | 62.63 | 83 | 0 | WP_323199853.1 |
| **Floridanemamide C (BLCC F167)** | | | | | | | | | |
| ***fmc*A** | Loading | 13,557 | A, KR, ACP | α-ketoisocaproic acid | *Microseira wollei* | 71.06 | 99 | 0 | WP_226584072.1 |
| | 1 | | KS, AT, KR, ACP | malonyl CoA | | | | | |
| | 2 | | KS, AT, KR, ACP | malonyl CoA | | | | | |
| ***fmc*B** | 1 | 11,061 | KS, AT, AmT, MO | malonyl CoA | *Desmonostoc muscorum* CCALA 125 | 69.05 | 100 | 0 | UBH04462.1 |
| | 2 | | C, A, E, PCP | D-Val | | | | | |
| ***fmc*C** | 1 | 25,200 | C, A, PCP | Gly | *Desmonostoc muscorum* LEGE 12446 | 66.74 | 100 | 0 | WP_255264359.1 |
| | 2 | | C, A, PCP | L-Gln | | | | | |
| | 3 | | C, A, PCP | L-Val | | | | | |
| | 4 | | C, A, E, PCP | D-*allo*-Ile | | | | | |
| | 5 | | C, A, PCP | L-Ser | | | | | |
| | 6 | | C, A, PCP | D-Leu | | | | | |
| | 7 | | C, A, PCP | L-Thr | | | | | |
| ***fmc*D** | 1 | 9,795 | C, A, E, oMT, PCP | D-hTyr | *Desmonostoc muscorum* LEGE 12446 | 67.35 | 89 | 0 | WP_255264360.1 |
| | 2 | | C, A, PCP | L-Pro | | | | | |
| ***fmc*E** | 1 | 4,188 | C, A, PCP, TE | L-Pro | Unclassified *Microcystis* (*IN: CYANOBACTERIA)* | 64.77 | 70 | 0 | WP_287686773.1 |

Table S5. Residue composition of cyanopeptides with beta amino acids

| **Compound Name** | **β-aa** | **1** | **2** | **3** | **4** | **5** | **6** | **7** | **8** | **9** | **10** | **Post-assembly line tailoring** |
|---|---|---|---|---|---|---|---|---|---|---|---|---|
| Tychonamide | Atpoa | D-Gln | Gly | Pro | Pro | D-*allo*-Ile | Ser | Dhb | Thr | D-O-Me-Htyr | Pro | N-Ac-N-Me-Leu |
| Floridanemamide A | Athmu | D-Gln | Gly | Ser | Phe | D-*allo*-Ile | Ser | Dhb | Thr | D-O-Me-Htyr | Pro | N-Butyl-Pro |
| Floridanemamide B | Athmu | D-Gln | Gly | Pro | Phe | Dhb | Ser | Dhb | Thr | D-O-Me-Htyr | Pro | N-Ac-N-Me-Leu |
| Pahayokolide A | Athmu | D-Gln | Gly | Pro | Phe | Dhb | Ser | Dhb | Thr | D-HPhe | Pro | N-Ac-N-Me-Leu |
| Portoamide A/Lyngbyazothrin C | Athmu | D-Gln | Gly | Pro | Pro | D-*allo*-Ile | Ser | Dhb | Thr | D-O-Me-Htyr | Pro | N-Ac-N-Me-Tyr |
| Floridanemamide C | Athmu | D-Val | Gly | Gln | Val | D-*allo*-Ile | Ser | D-Leu | Thr | D-O-Me-Htyr | Pro | N-Butyl-Pro |
| Schizotrin A | Athmu | D-Gln | Gly | Pro | Phe | D-Val | Ser | Dhb | Ser | O-Me-Htyr | Pro | N-Butyl-N-Me-D-Ala |
| Scytonemin A | Ahda | D-Ser | Gly | HyMePro | HyMePro | D-Leu | Hser | D-Phe | Gly | D-HyLeu | MePro | N-Ac-Ala |
| Muscotoxin A | Ahdoa | D-Gln | Gly | Pro | Phe | D-*allo*-Ile | Ser | Dhb | Ser | D-*allo*-Ile | Pro | |

Table S6. Genomic information for analyzed cyanobacterial strains of *Floridanema*

| Strain | *F. flaviceps* BLCC-F50 | *F. evergladense* BLCC-F167 | *F. aerugineum* BLCC-F306 |
|---|---|---|---|
| Completion (%) | 99.94 | 100 | 100 |
| Contamination (%) | 0.1 | 0.05 | 0.06 |
| Coverage Illumina (x) | 29.48 | 19.6 | 124.1 |
| Coverage ONT (x) | 88.8 | 60.1 | 84.8 |
| Contigs | 3 | 1 | 12 |
| Genome Size (bp) | 6750723 | 6463832 | 6967955 |
| GC % | 40.7 | 40.1 | 40.5 |
| tRNA | 73 | 75 | 76 |
| rRNA | 9 | 9 | 9 |
| CDS | 6196 | 5981 | 6234 |
| SRA | SRS22629764 | SRS22630153 | SRS29217320 |
| BioSample | SAMN43549591 | SAMN43549593 | SAMN60208900 |

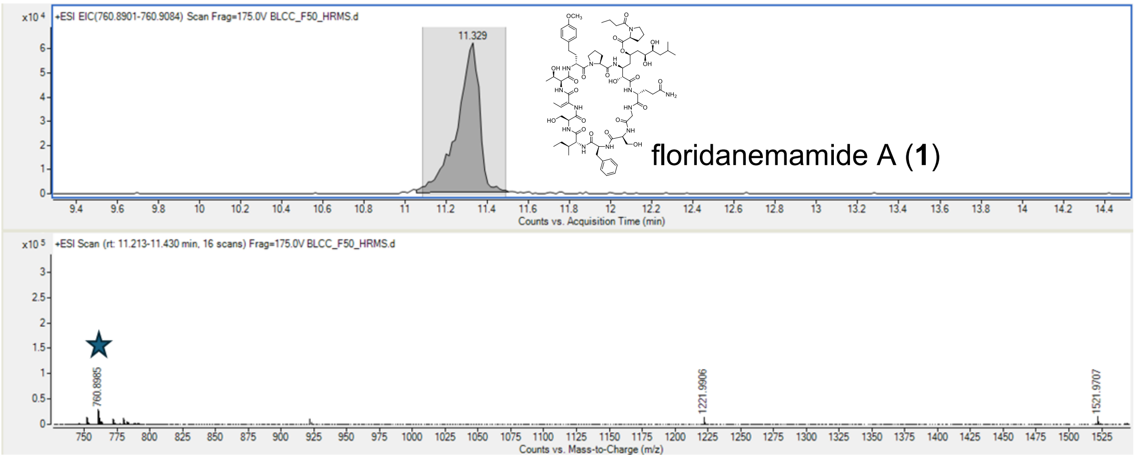


Figure S1. HRESIMS measurement of floridanemamide A (**1**) *m/z* 760.8985 $[M+2H]^{2+}$.

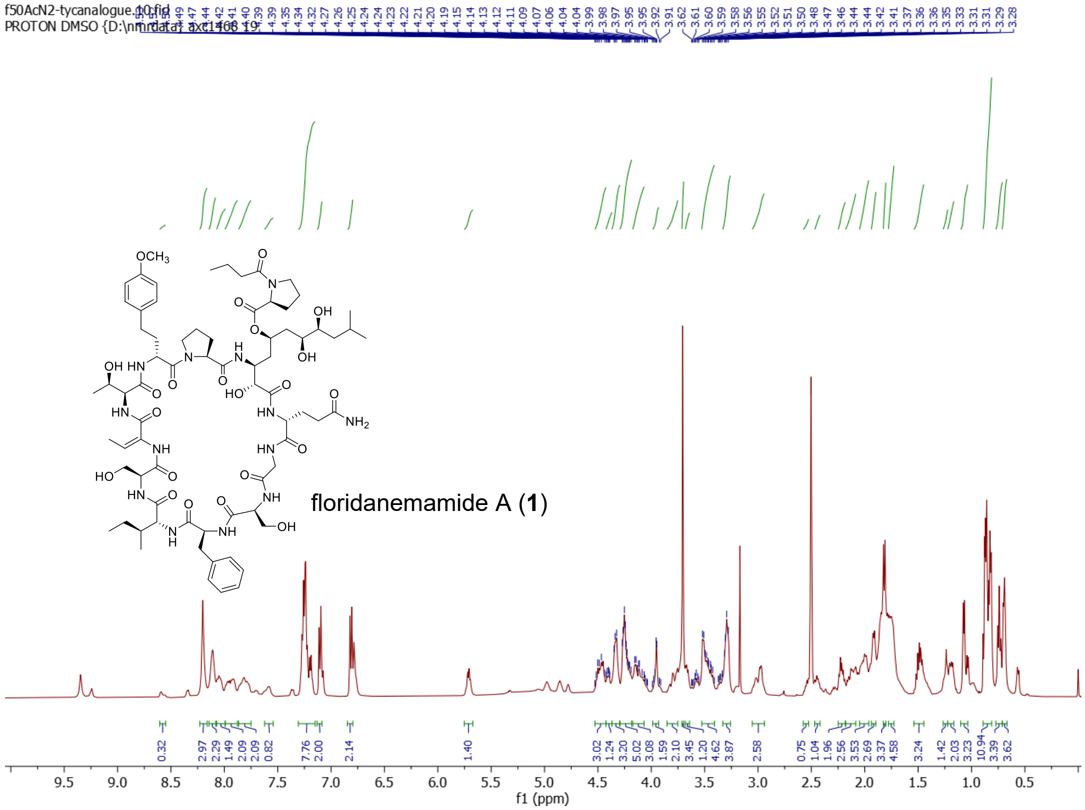


Figure S2. $^{1}$H NMR of floridanemamide A (**1**) (500 MHz, DMSO-$d_6$).

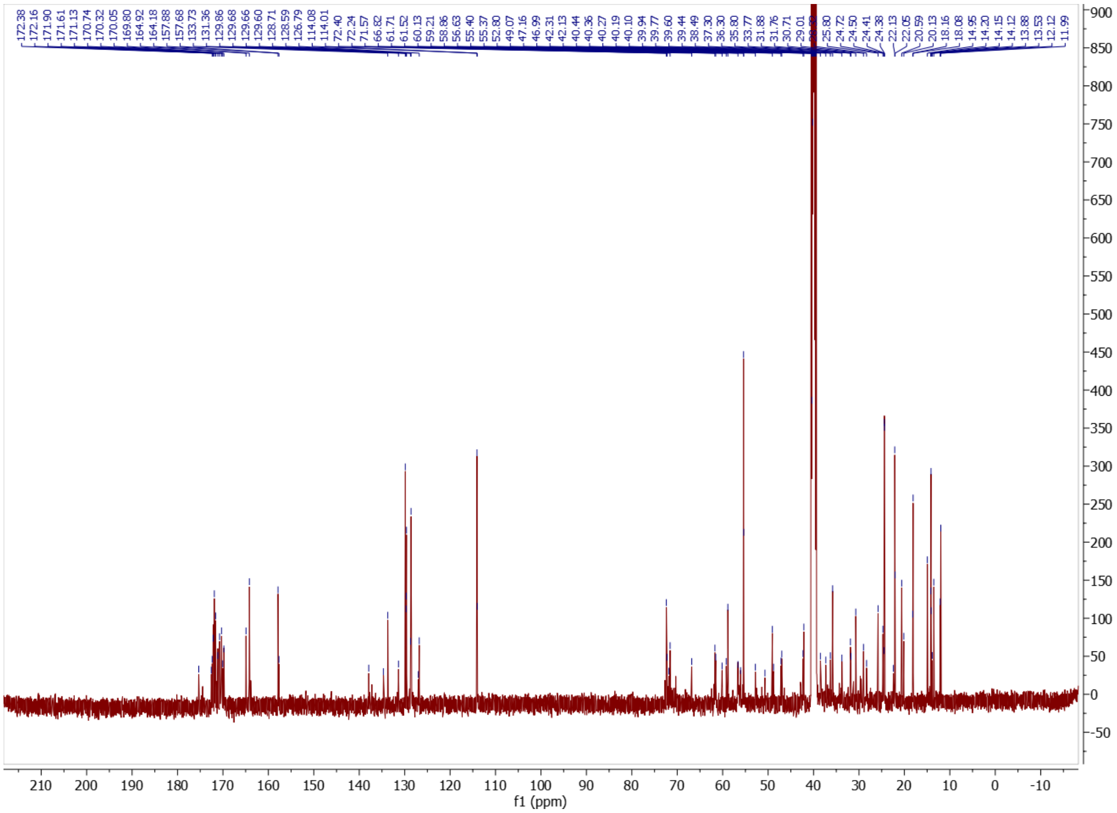


Figure S3. $^{13}C$ NMR of floridanemamide A (125 MHz, DMSO-$d_6$).

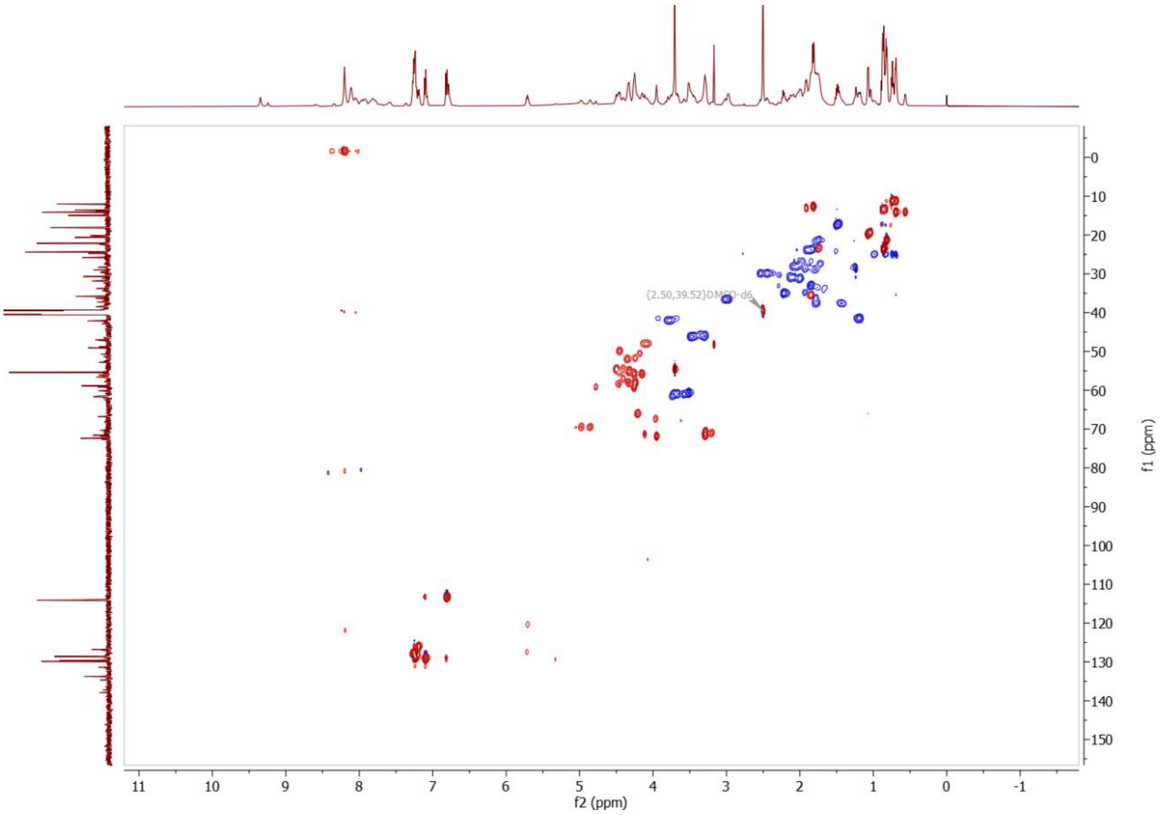


Figure S4. Multiplicity-edited HSQC of floridanemamide A.

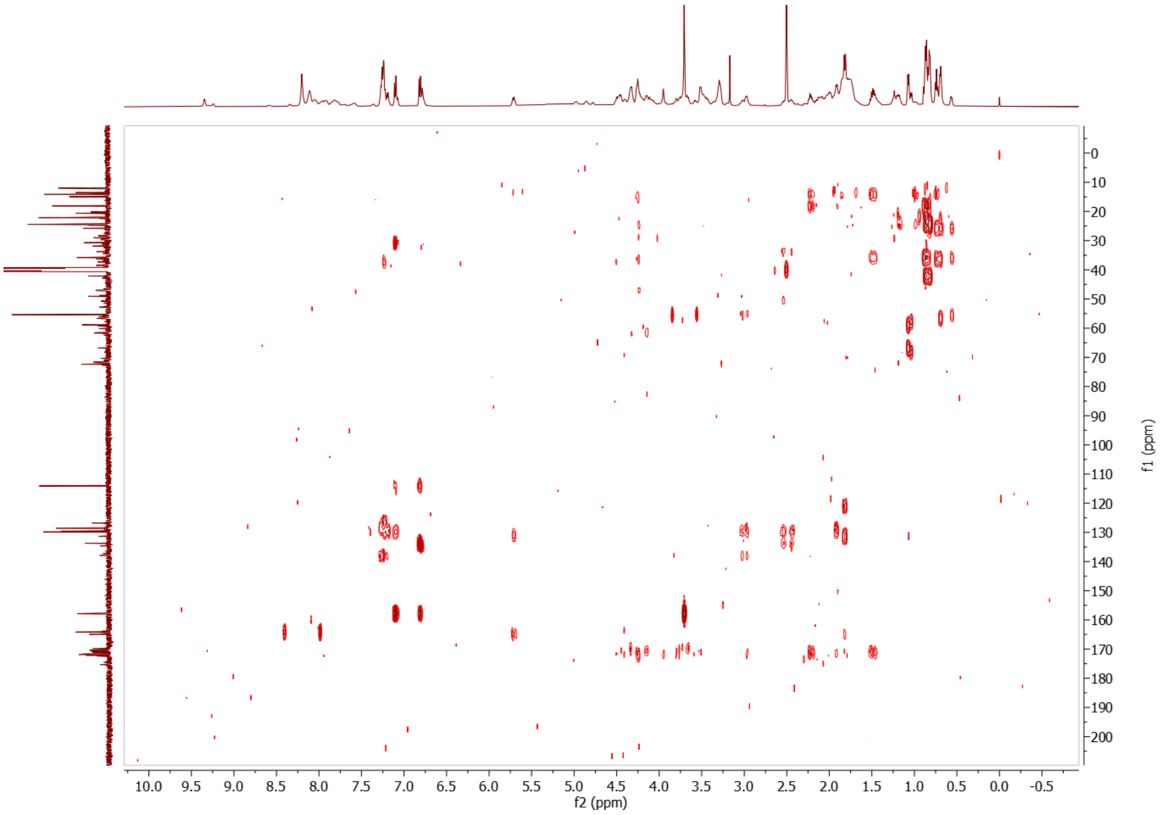


Figure S5. HMBC of floridanemamide A.

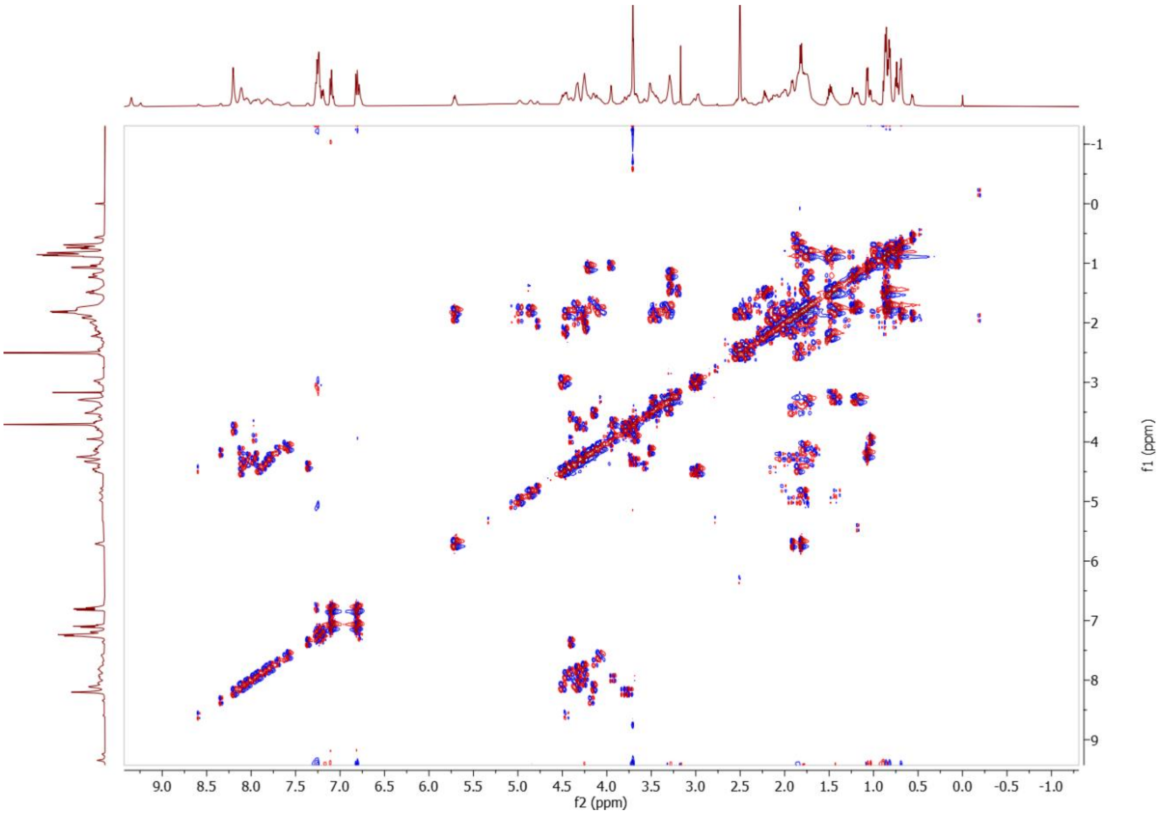


Figure S6. DQF-COSY of floridanemamide A.

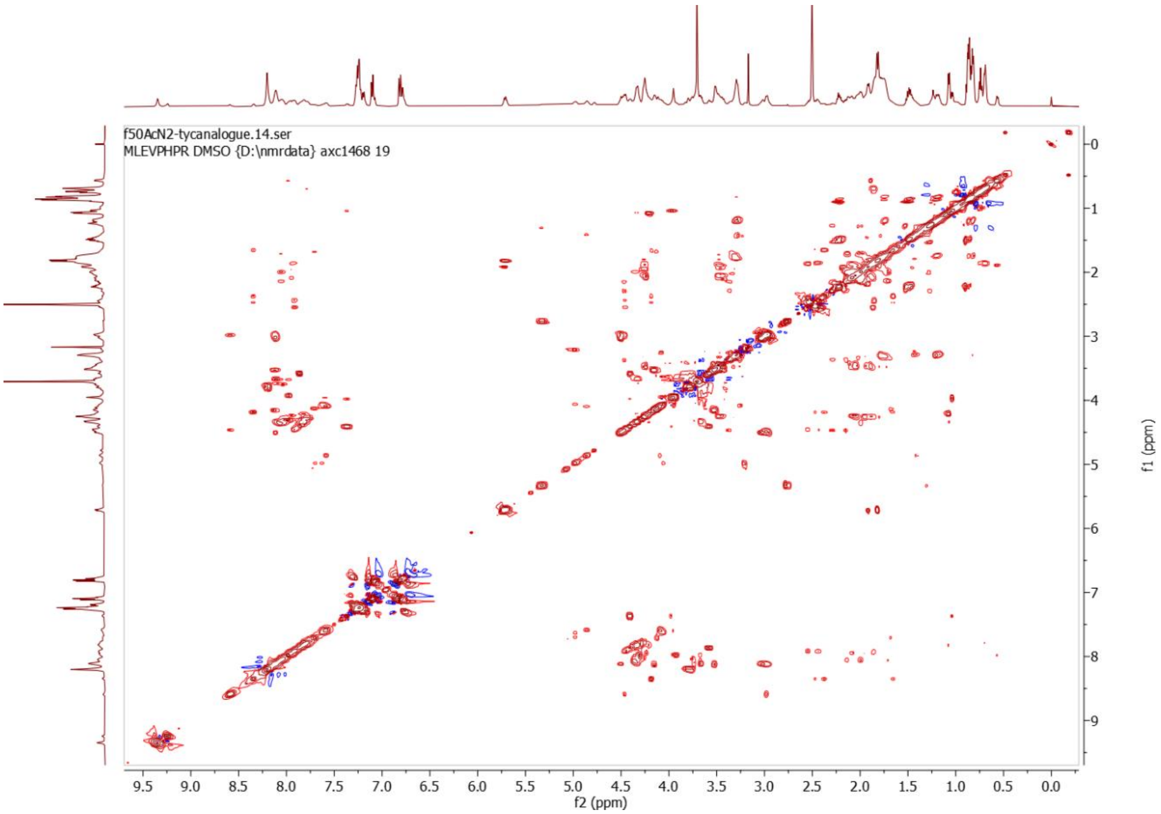

Figure S7. TOCSY of floridanemamide A.

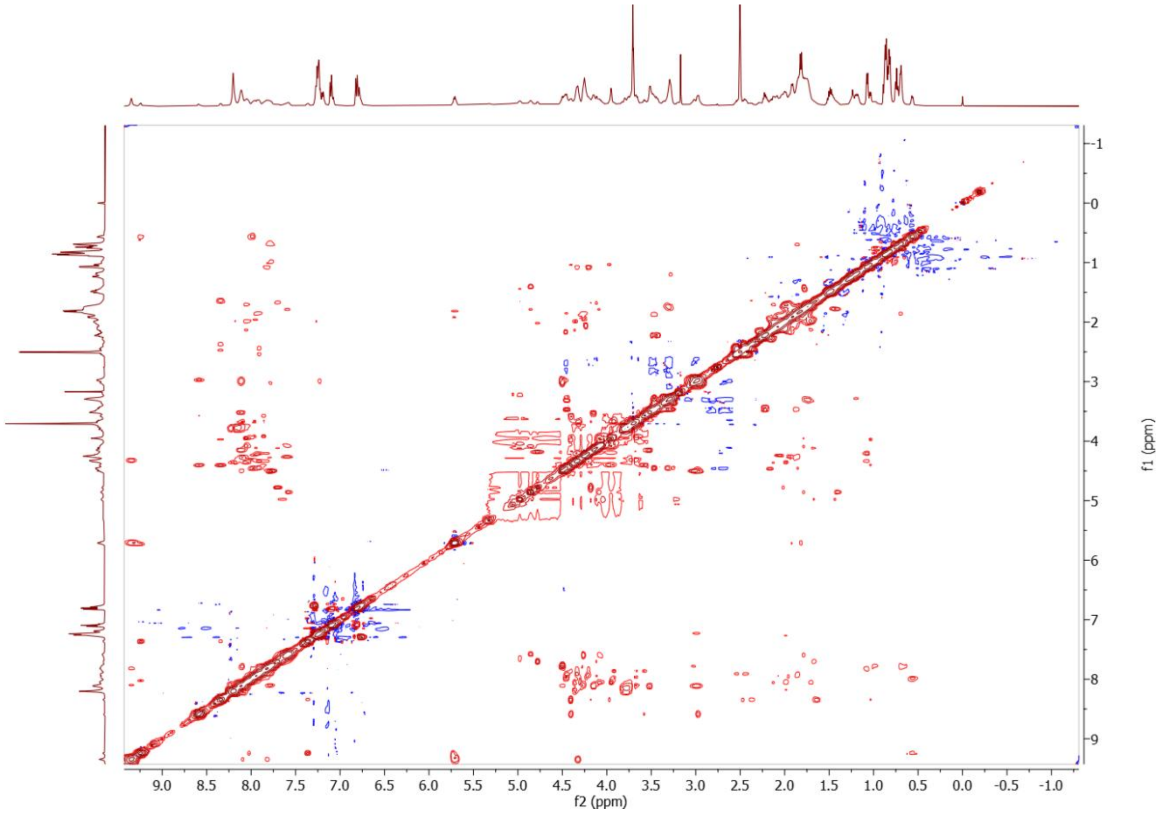


Figure S8. NOESY of floridanemamide A.

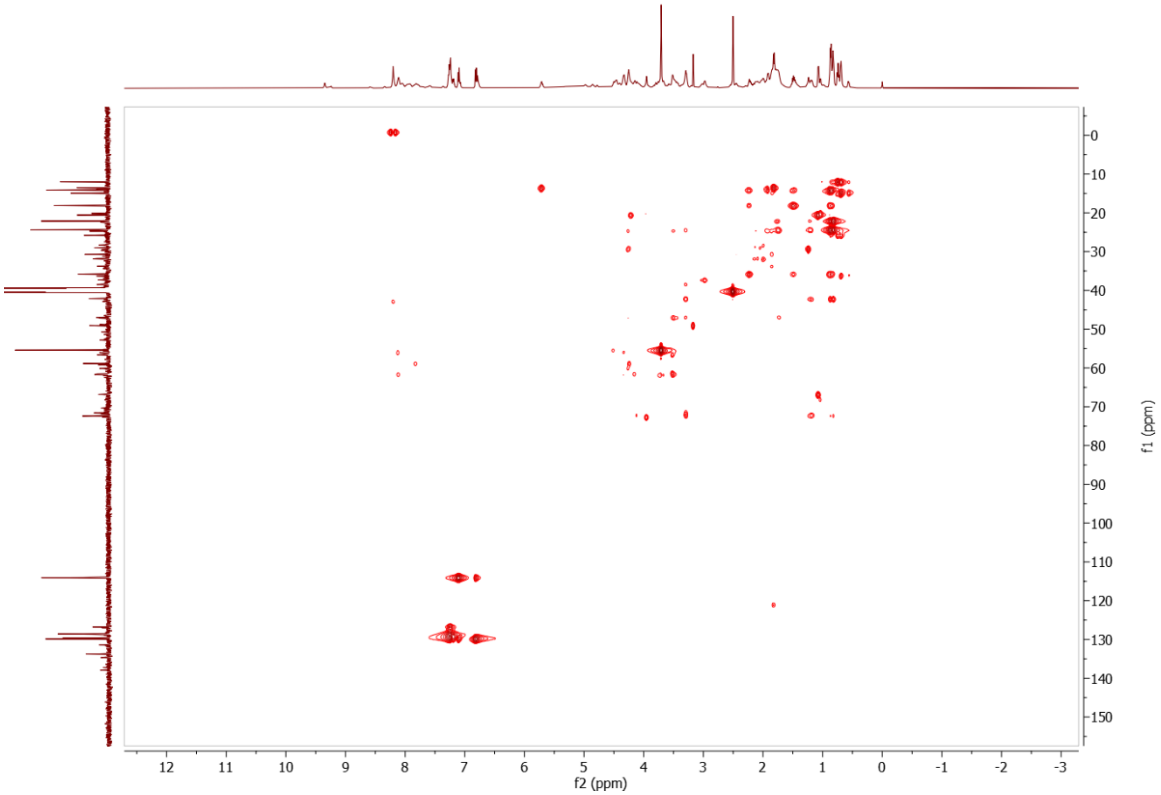


Figure S9. HSQC-TOCSY of floridanemamide A.

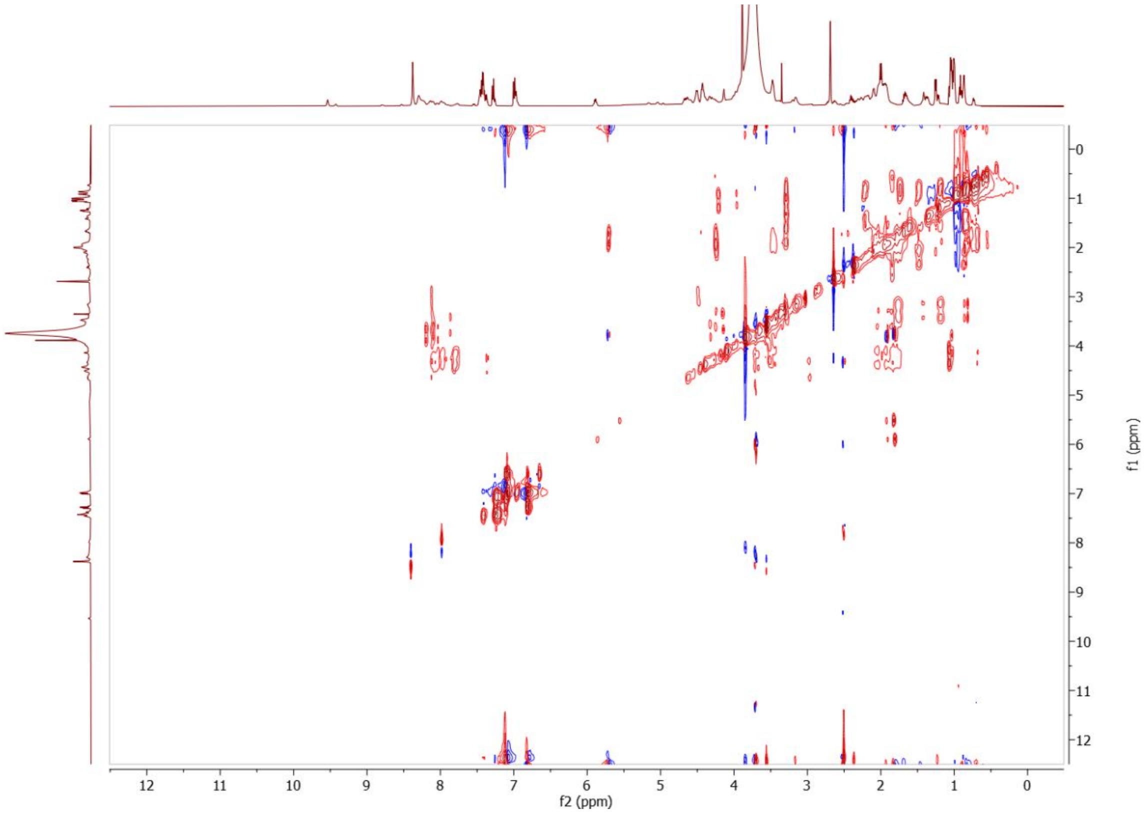

Figure S10. HETLOC of floridanemamide A.

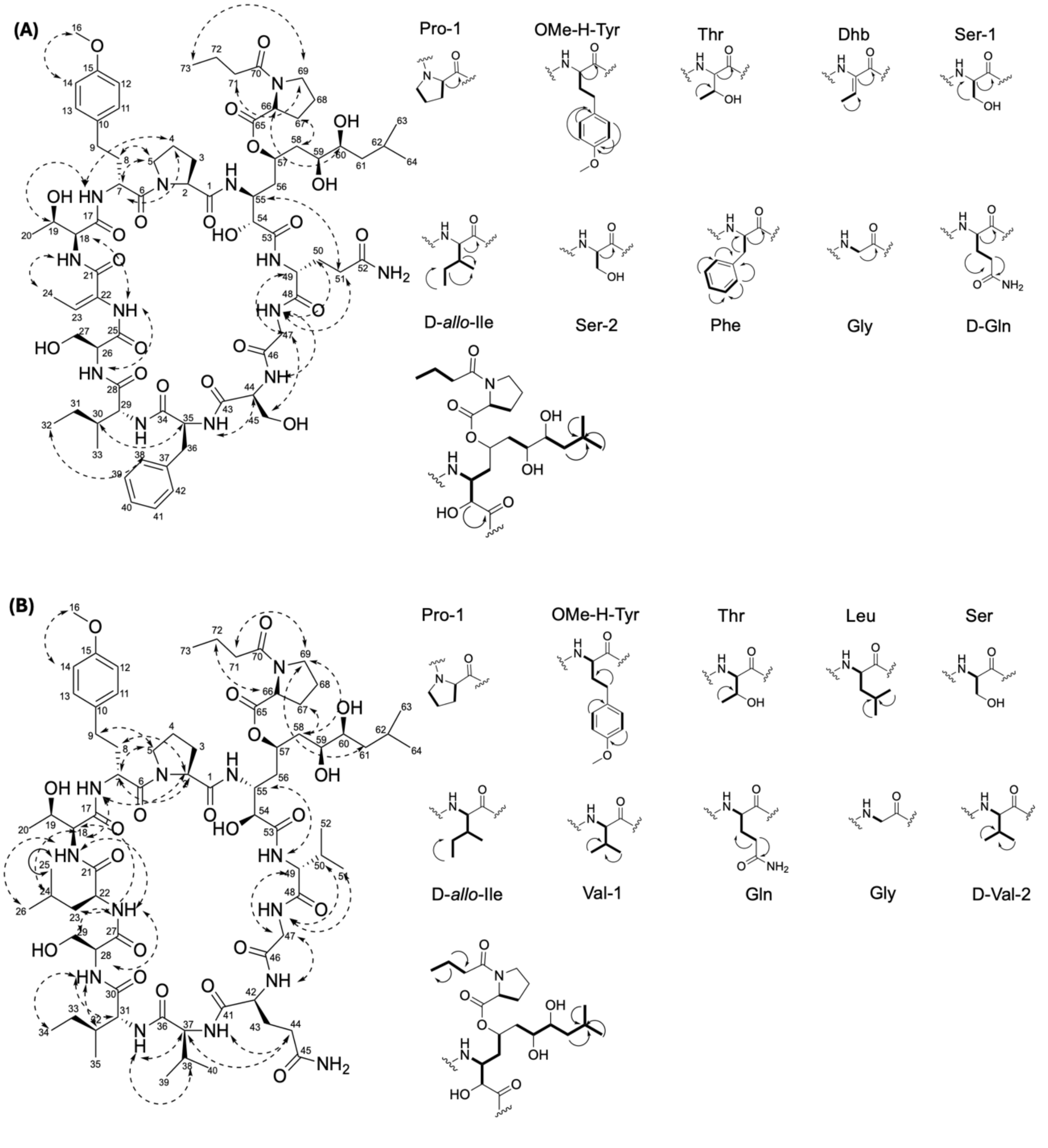


Figure S11. Select 2D NMR correlations for (A) floridanemamide A (**1**) and (B) floridanemamide C (**3**). TOCSY correlations shown with bold lines, HMBC correlations shown with arrows and NOE correlations shown with dashed arrows.

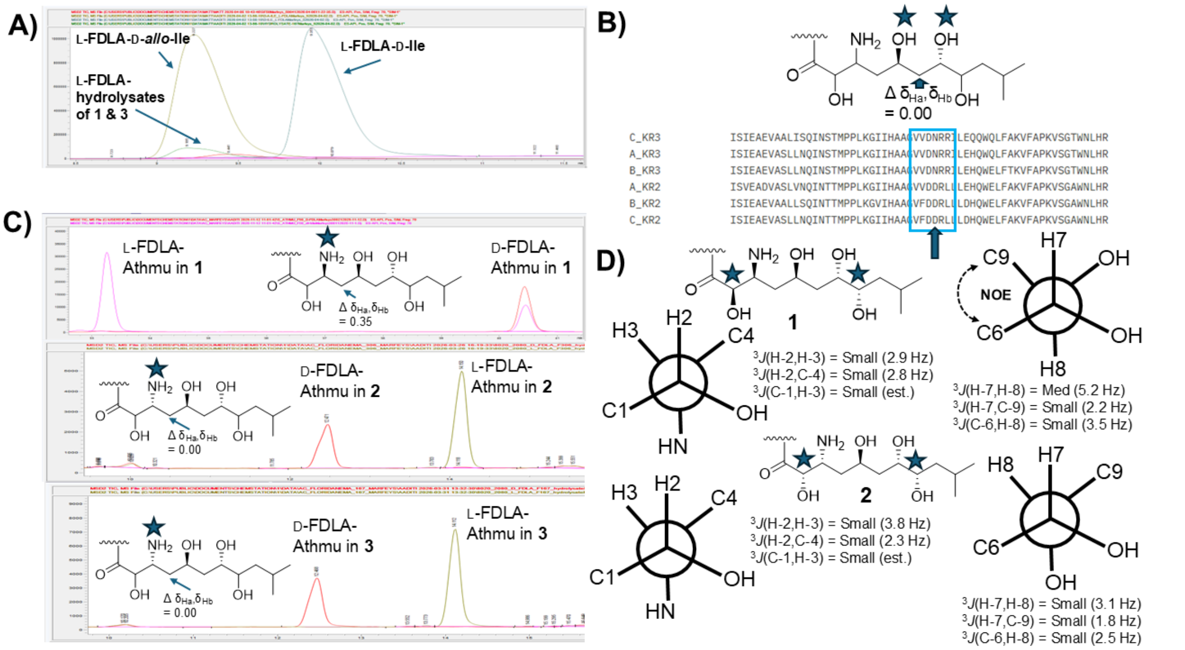


Figure S12. Relative and absolute configuration analysis of **1-3**. (A) Marfey's analysis of the hydrolysates of **1** and **3** reacted with L-FDLA and compared to L-FDLA reacted standards. (B) Sequence analysis of the KR domains predicted to reduce β-ketones in the *fma* pathway with the VDD sequence indicative of B-type KRs and the VDN sequence indicative of A-type KRs noted in the blue box. Additionally, the difference between the two protons attached to the intervening methylene unit is noted, which was 0.00 for **1-3** (blue arrow). (C) The determination of C-3 configuration of the Athmu unit by reacting the hydrolysates of **1-3** with L- and D- FDLA, respectively. Additionally, the difference between the two protons attached to the intervening methylene unit between C-3 and C-5 is noted for **1-3**, which was 0.35 for **1** and 0.00 for **2** and **3**. (D) *J*-coupling analysis of portions the Athmu unit to determine the relative configuration between C2 and C3 and C7 and C8 in **1** and **2**. The coupling constants for **3** exhibited medium (4.6 Hz), small (2.5 Hz), and small (est.) respective *J* values for H2-H3, H2-C4 and H3-C1 and small, small, small values for H7-H8, H7-C9, and H8-C6, respectively.

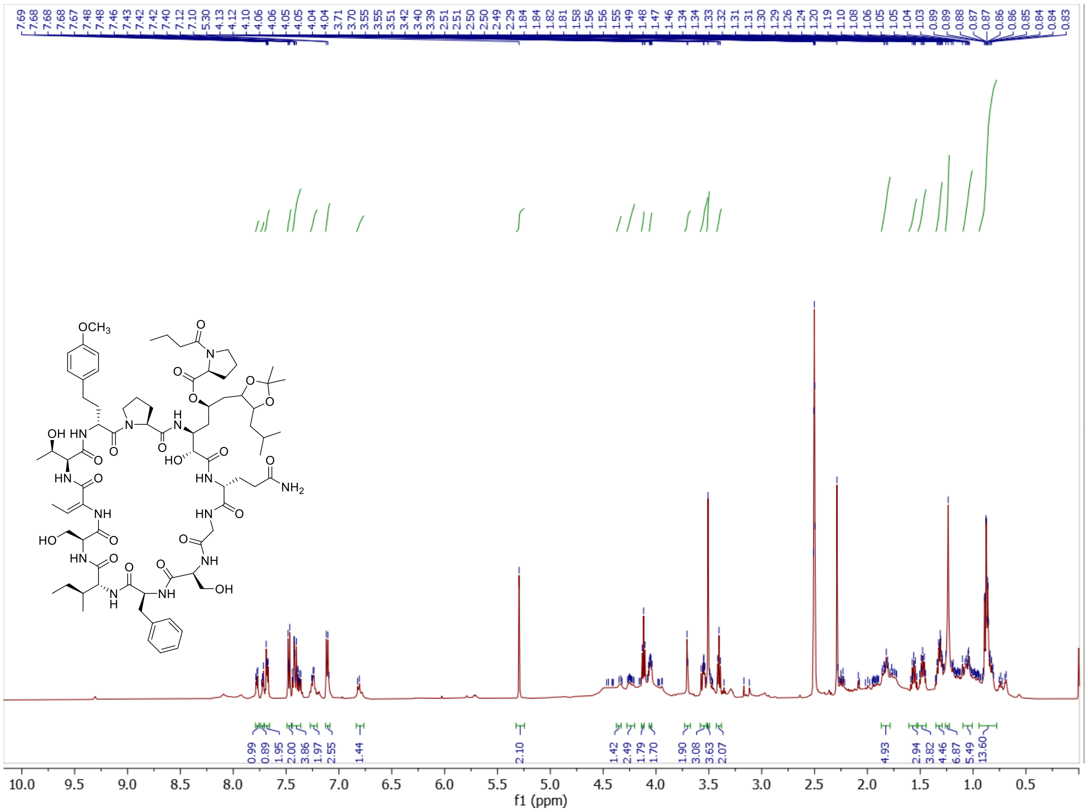

Figure S13. [1]H NMR of the floridanemamide A acetonide derivative (500 MHz, DMSO-$d_6$).

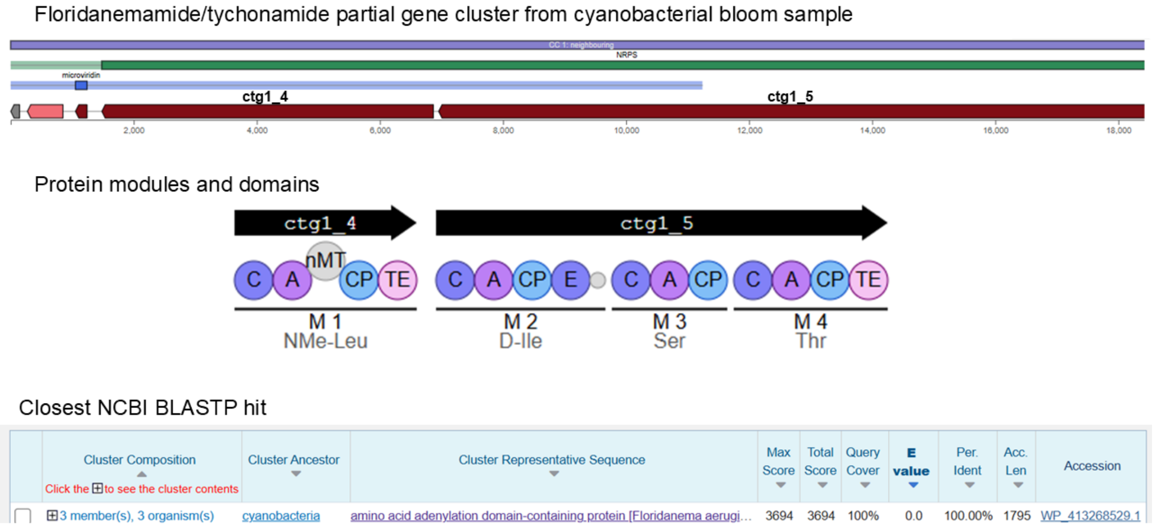


Figure S14. Partial antiSMASH (version 8) annotation of partial floridanemamide/tychonamide pathway in Lake Erie, Showse Park MAG. Protein modules and domains are shown below the open reading frames and the closest BLASTP hit for contig1_4 was *Floridanema aerugineum* with 100% amino acid identity with a 1795 accession length.

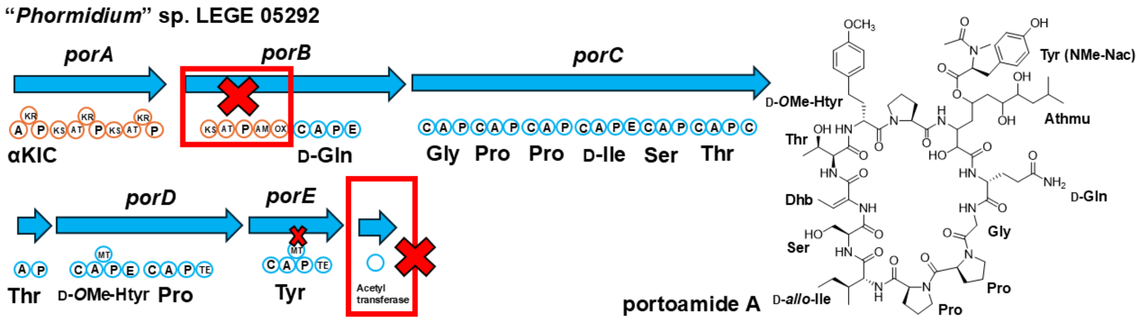


Figure S15. Putative portoamide (*por*) biosynthetic pathway with the portoamide A structure shown at right. Red boxes show portions of pathway that were not annotated but are predicted for biosynthesis. For abbreviations, see Figure 1 legend.

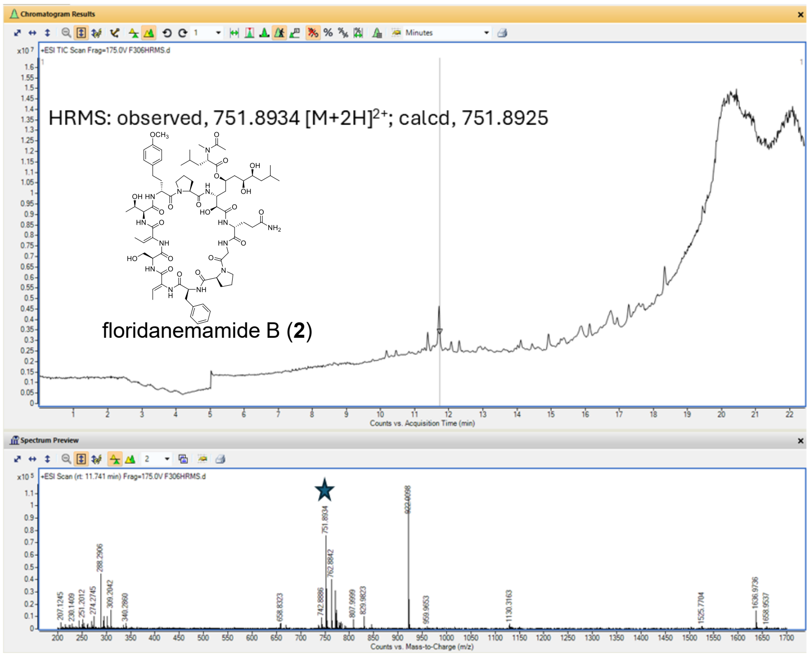


Figure S16. HRESIMS measurement of floridanemamide B (**2**) *m/z* 751.8934 $[M+2H]^{2+}$.

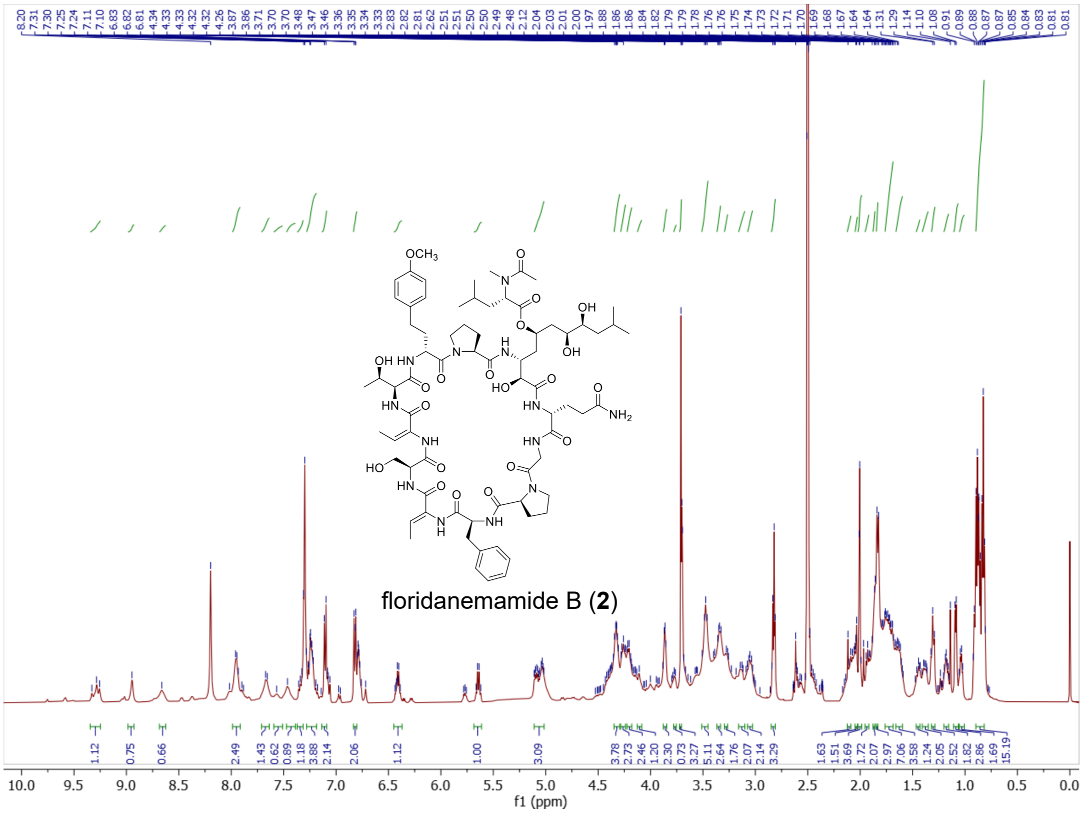


Figure S17. $^{1}$H NMR of floridanemamide B (**2**) (500 MHz, DMSO-$d_6$).

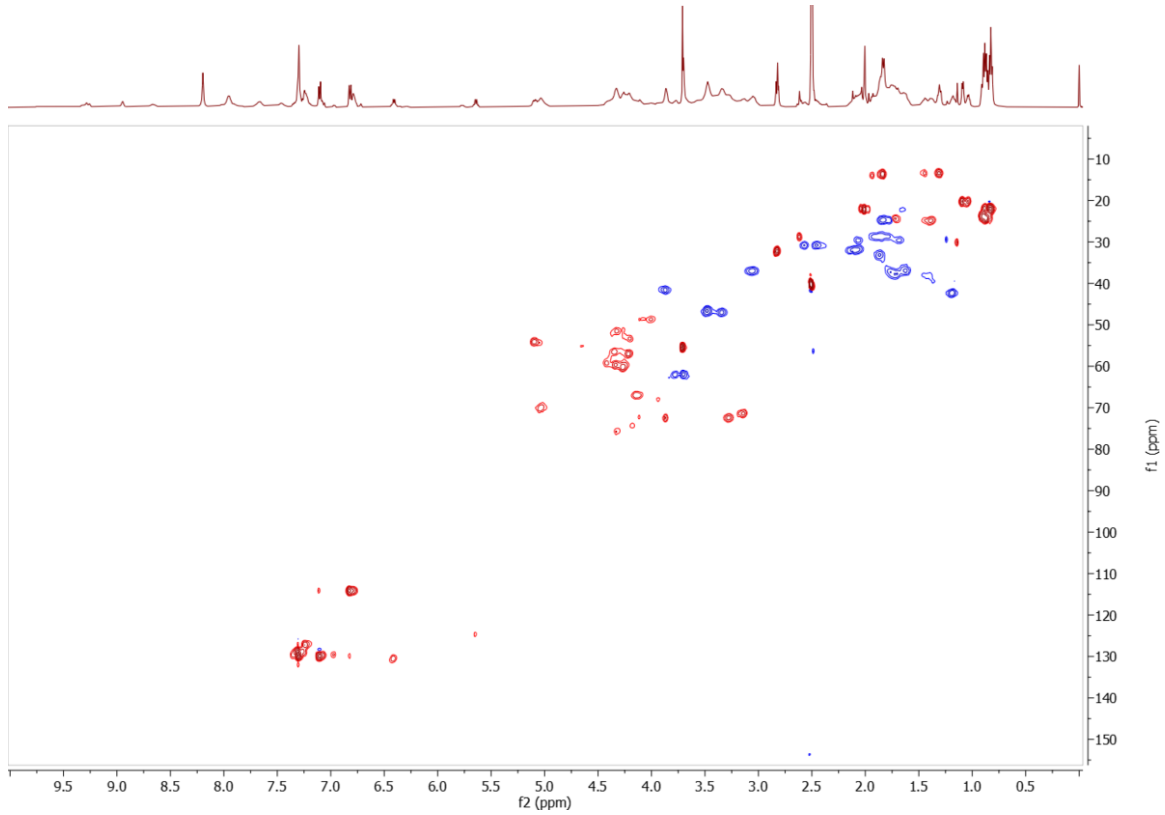


Figure S18. Multiplicity-edited HSQC of floridanemamide B.

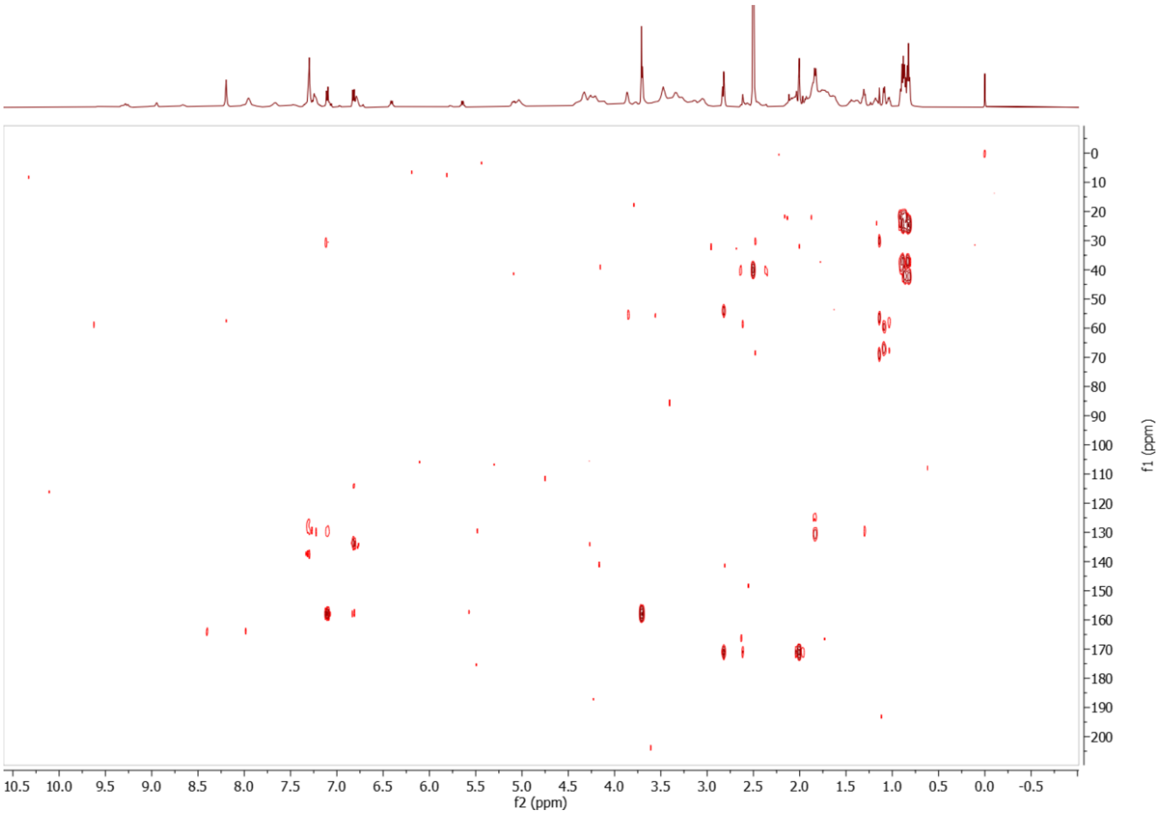


Figure S19. HMBC of floridanemamide B.

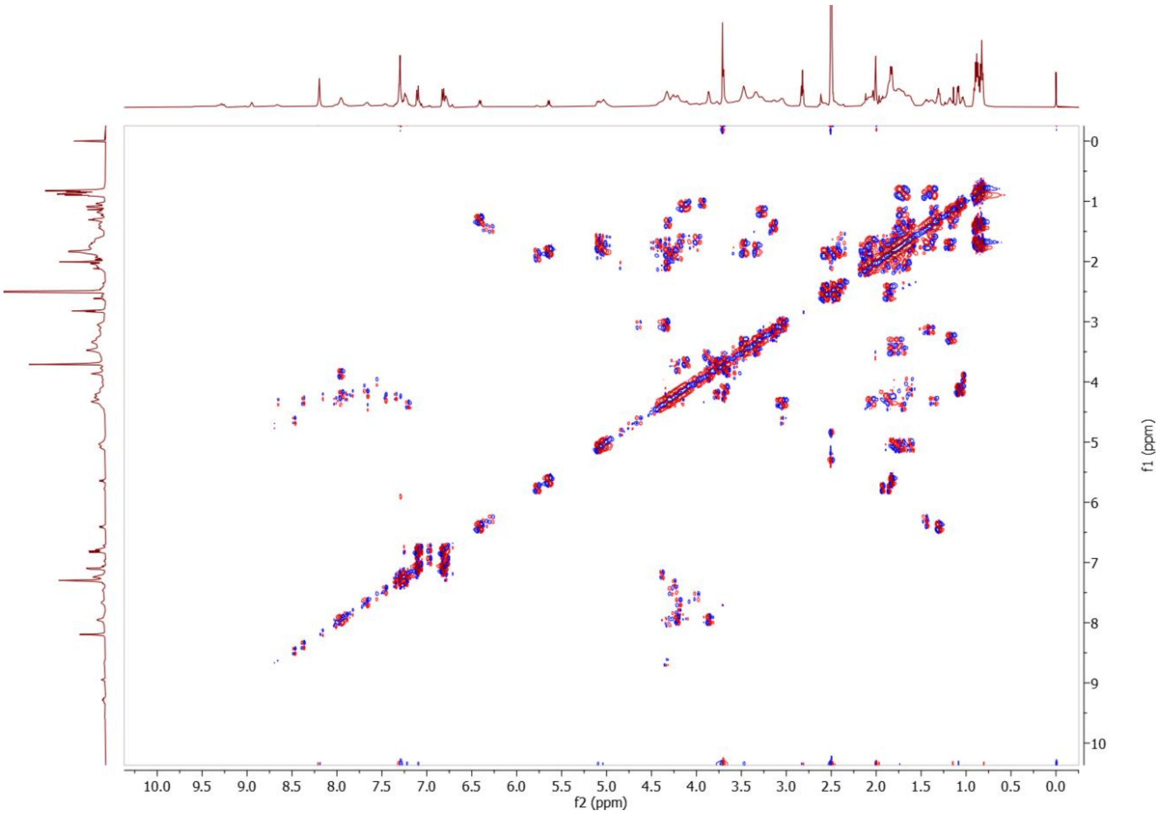


Figure S20. DQF-COSY of floridanemamide B.

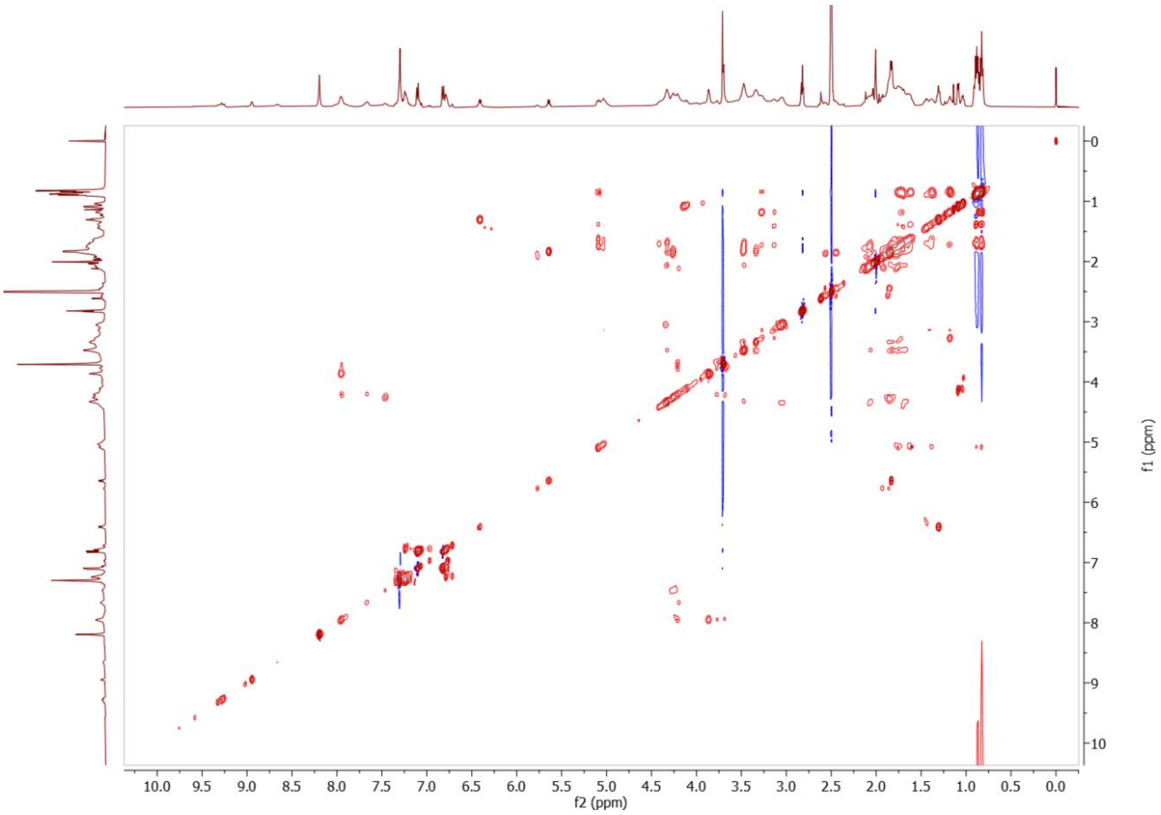


Figure S21. TOCSY of floridanemamide B.

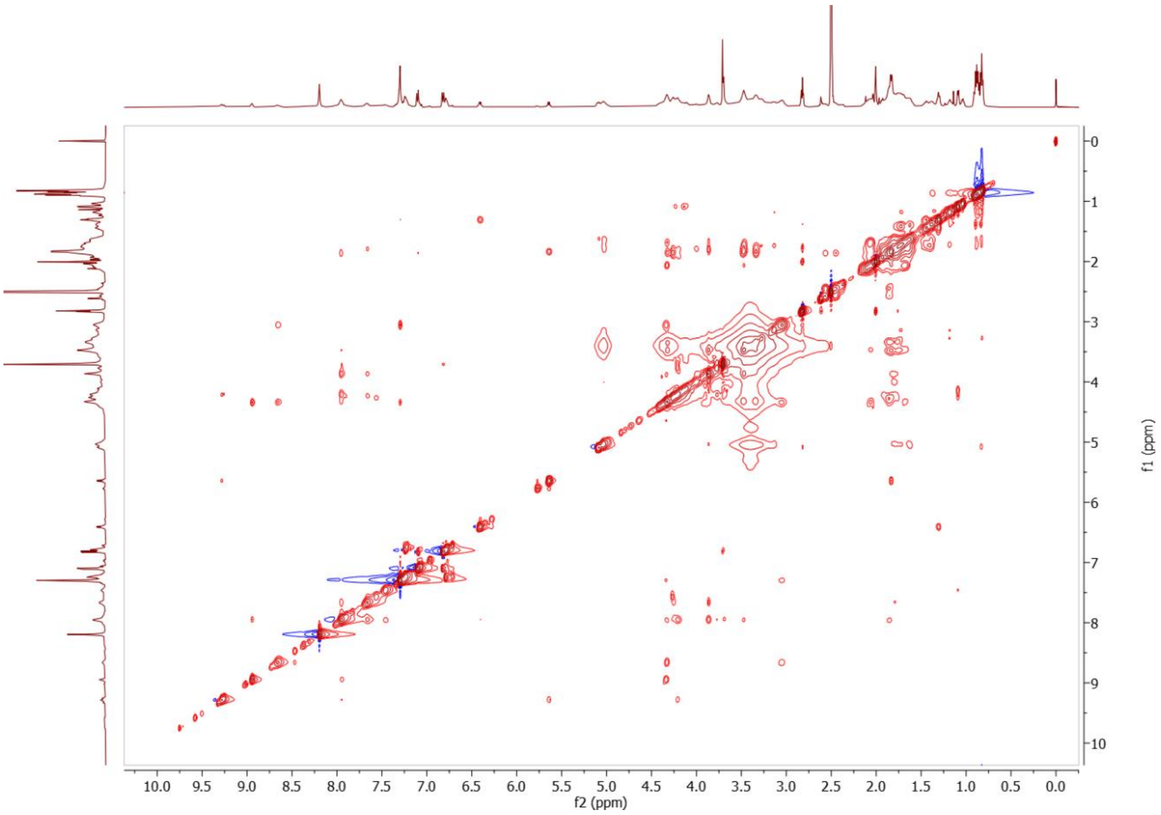


Figure S22. NOESY of floridanemamide B.

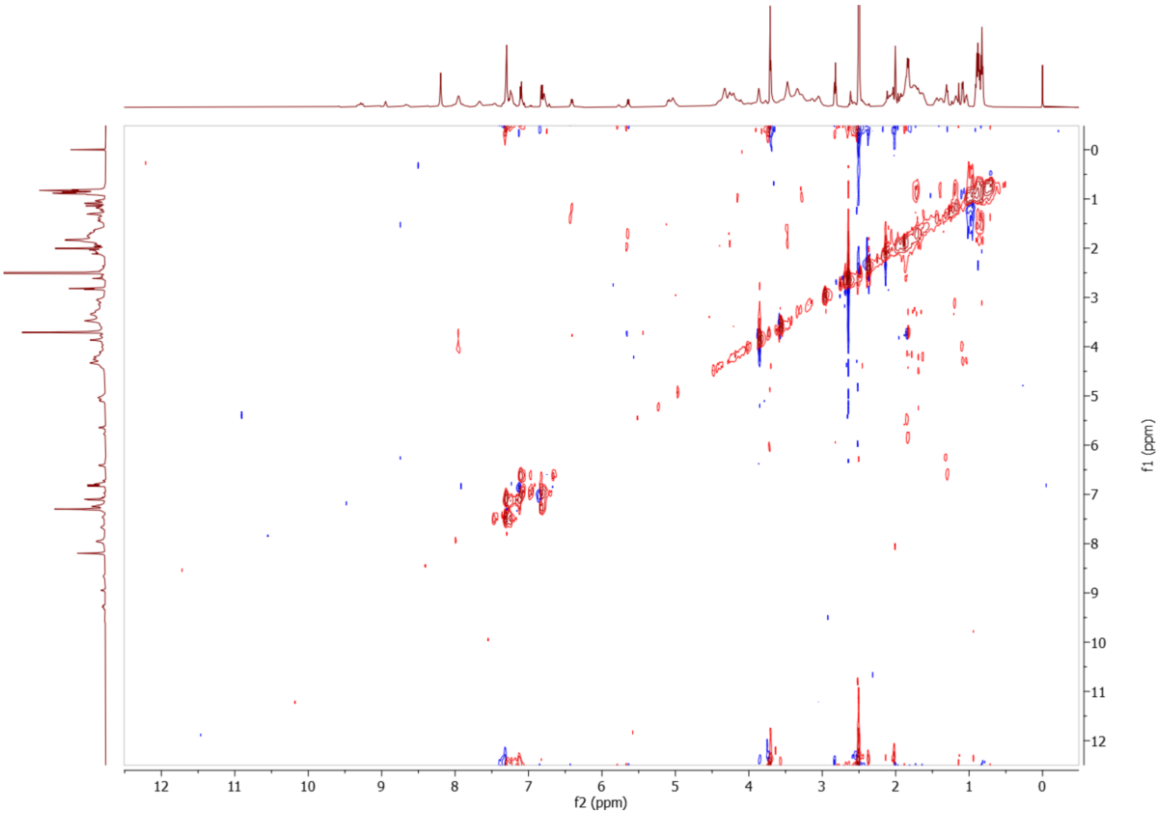


Figure S23. HETLOC of floridanemamide B.

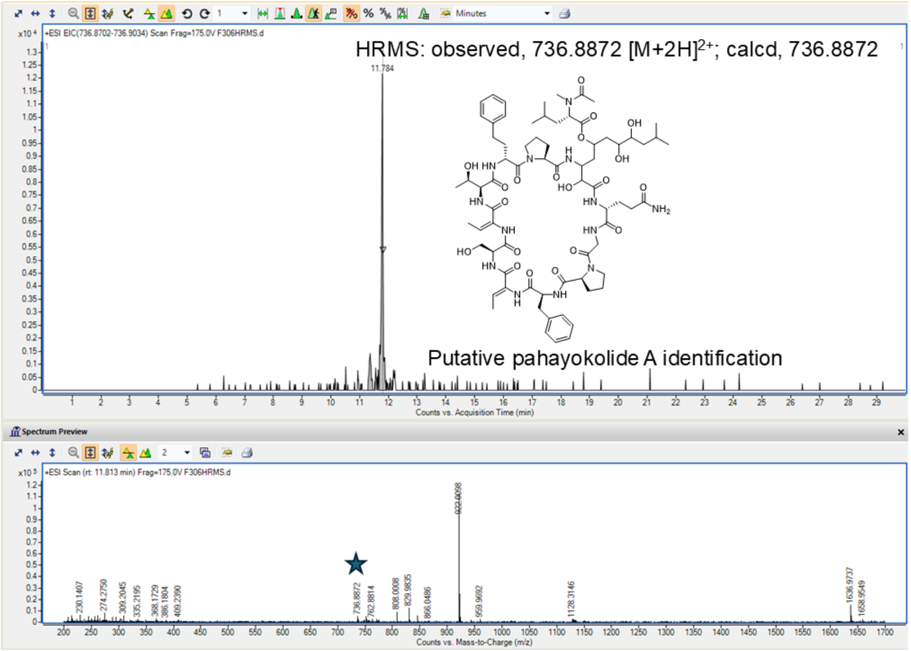


Figure S24. HRESIMS measurement of putative pahayokolide A *m/z* 736.8872 $[M+2H]^{2+}$.

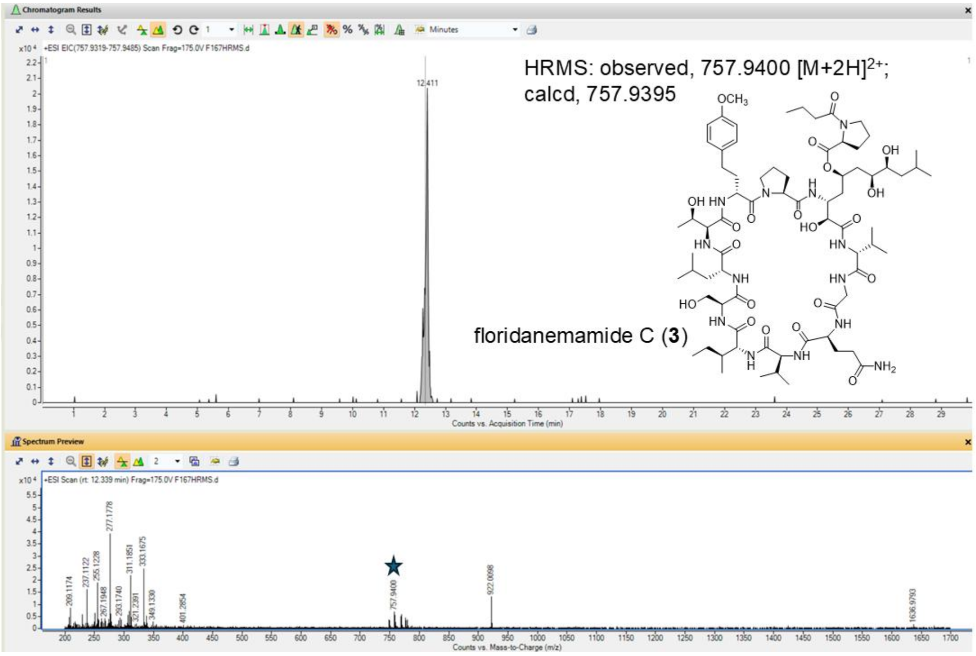


Figure S25. HRESIMS measurement of floridenemamide C (**3**) *m/z* 757.9400 $[M+2H]^{2+}$.

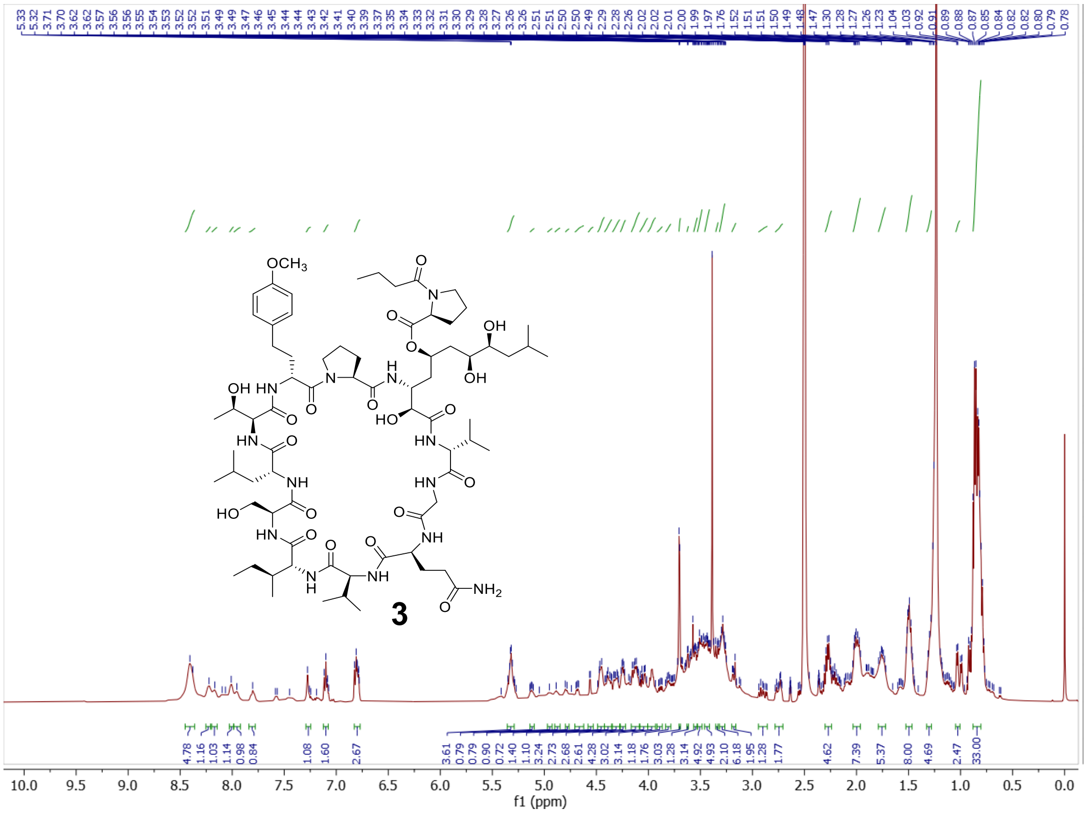

Figure S26. $^1$H NMR of floridanemamide C (**3**) (500 MHz, DMSO-$d_6$).

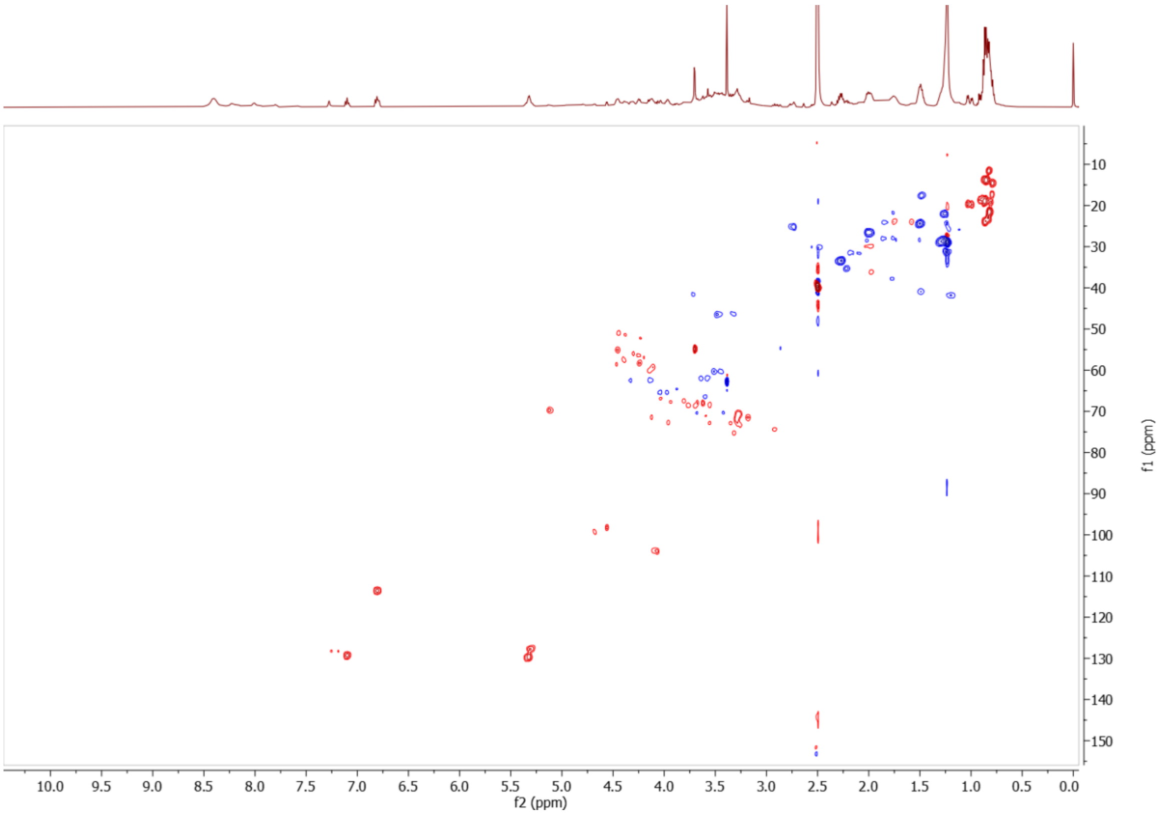


Figure S27. Multiplicity-edited HSQC of floridanemamide C.

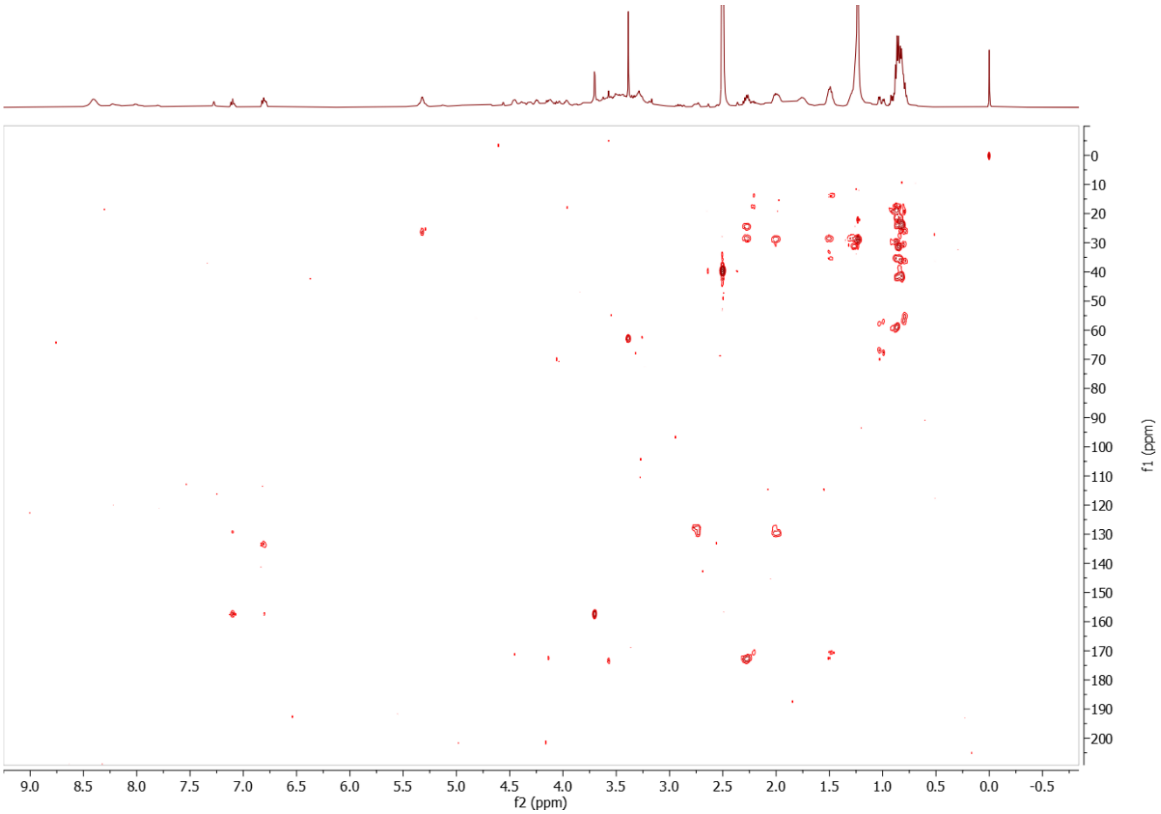

Figure S28. HMBC of floridanemamide C.

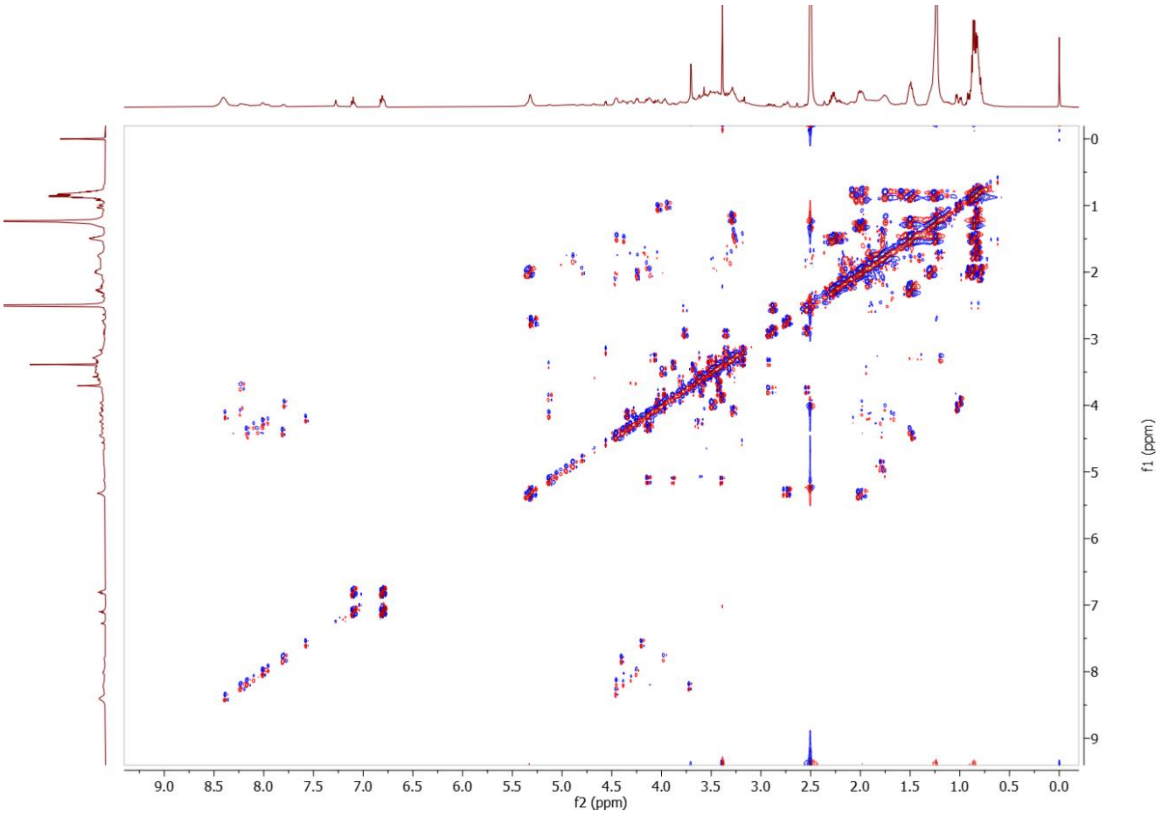


Figure S29. DQF-COSY of floridanemamide C.

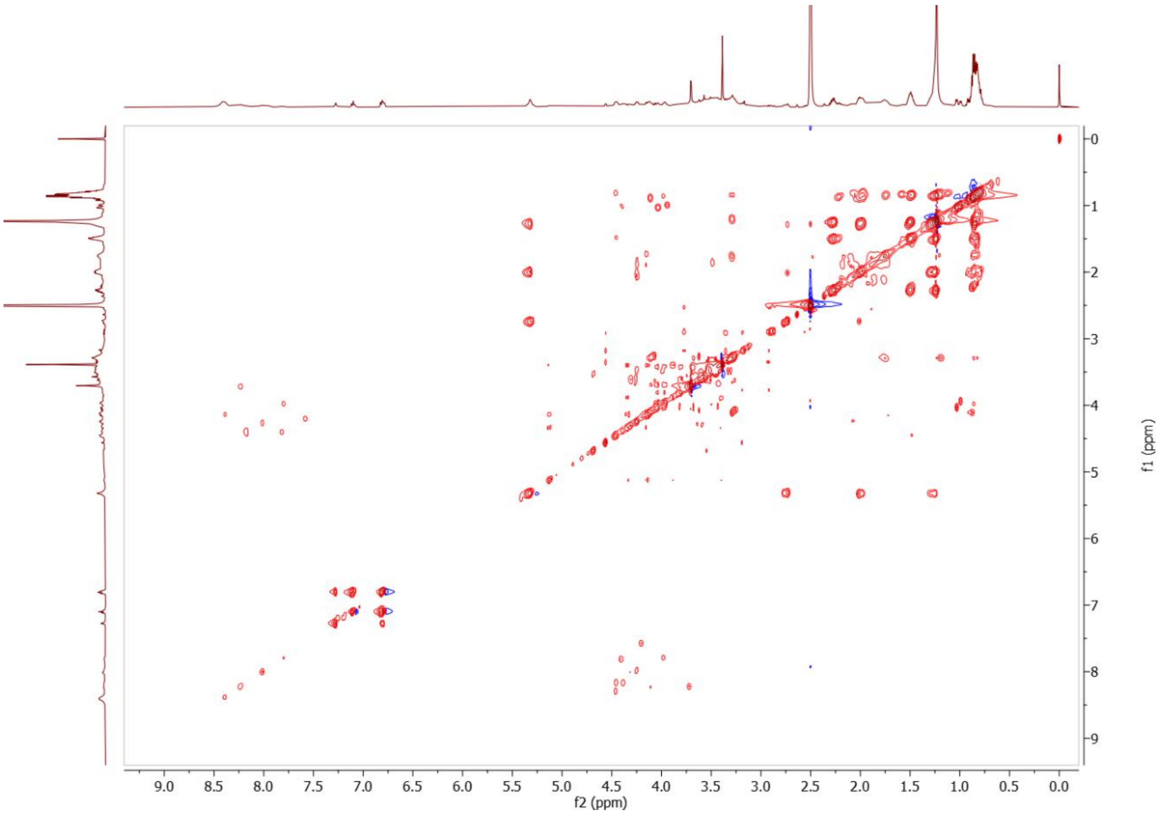

Figure S30. TOCSY of floridanemamide C.

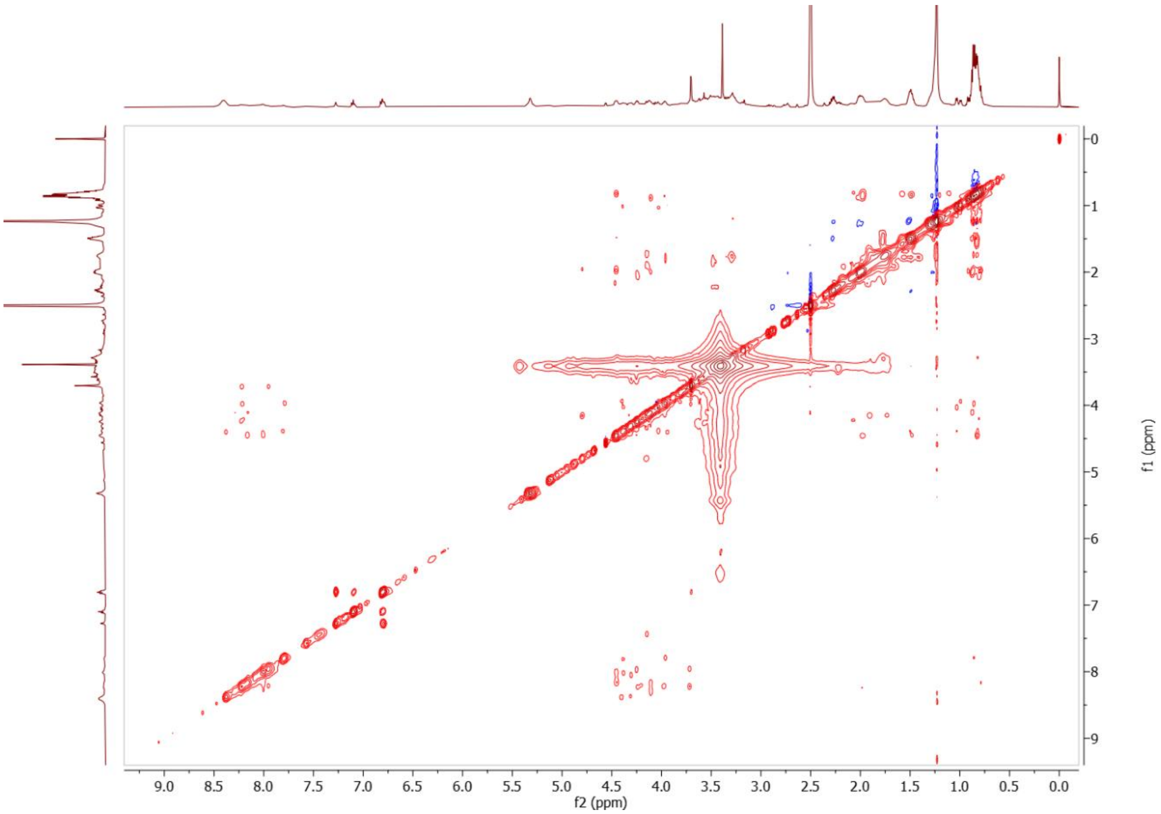

Figure S31. NOESY of floridanemamide C.

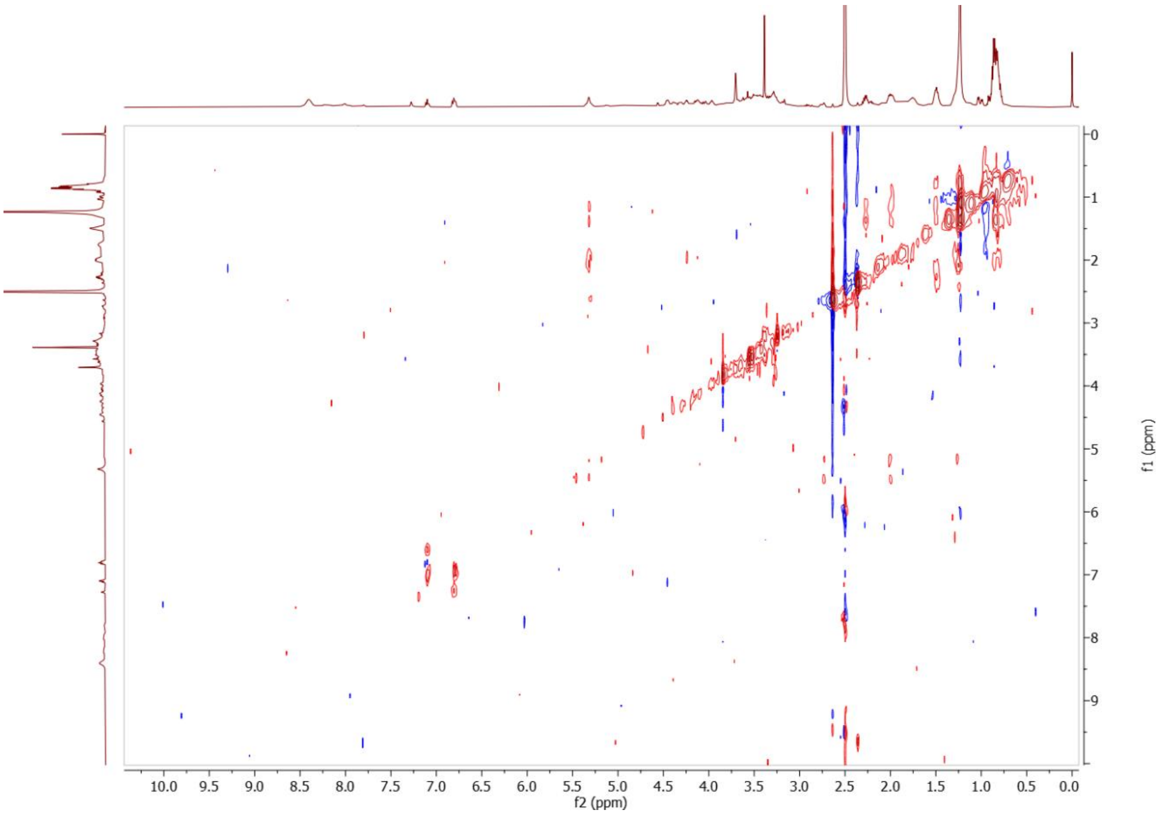

Figure S32. HETLOC of floridanemamide C.
